# Reference-free compound identification using computational prediction of molecular properties and multi-dimensional spectrometric measurements: a fentanyl case study

**DOI:** 10.64898/2026.04.22.719980

**Authors:** Christopher P. Harrilal, Adam L. Hollerbach, Danielle Ciesielski, Katherine J. Schultz, Richard Overstreet, Peter S. Rice, Ethan King, Julia Nguyen, Dylan H. Ross, Vivian S. Lin, Grace Y. Deng, Eva Brayfindley, Bobbie-Jo M. Webb-Robertson, Simone Raugei, Yehia M. Ibrahim, Robert G. Ewing, Thomas O. Metz

## Abstract

Mass spectrometry is used to identify chemicals to which humans are exposed, but it cannot directly determine molecular structures. Instead, structures are inferred by matching experimental spectra to libraries of spectra constructed from analyses of pure reference compounds. However, the chemical space of human exposures far exceeds the amount of experimental library spectra. Here, we evaluate a ‘reference-free’ strategy for confident identification of unknown molecules. Using fentanyl as a case study, we created a suspect library of over 1 billion computationally predicted fentanyl analogs and predicted molecular properties through machine learning, molecular dynamics, and density functional theory. Multi-dimensional spectra from a blinded analysis of a mock fentanyl tablet were matched with the predicted library, yielding an average of three candidate structures per measured analog, with six exact identifications. This work emphasizes the promise of reference-free molecular measurements for assessing human exposure by merging computational predictions with high-dimensional measurements.

## Main

Human development and long-term health are determined through continuous and dynamic interactions between the genome and the environment, historically considered as G x E interactions.^1-3^ The contribution of the environment – including the microbiome, chemicals, pathogens, and psychosocial and economic factors – was conceptualized as the ‘exposome’ by Wild in 2005,^4^ and its collective impact on human health is studied through exposomics.^5, 6^ Tobacco use^7^ and asbestos exposure^8^ are well-established components of the exposome that have resulted in key regulatory changes and global health improvements,^9, 10^ and a growing body of literature is now contributing to an increased understanding of the detrimental health effects of human exposure to per- and polyfluoroalkyl substances (PFAS)^11^. Continued urbanization, establishment of power and computing centers to support the growing AI market, and climate change are expected to increase the rate and types of human exposures. Today, several collections of environmentally-relevant chemicals are available to exposome researchers and the regulatory community, and highlight the chemical space that must be considered when conducting human exposomics studies: the Blood Exposome database, containing 67,291 xenobiotic chemicals reported to be found in human blood;^12^ PubChem Lite, which is a subset of 569,233 chemicals in PubChem relevant for exposomics studies;^13^ and the U.S. Environmental Protection Agency’s Distributed Structure-Searchable Toxicity (DSSTox) database, with over 1.2 million chemicals.^14^

Mass spectrometry (MS) has emerged as a workhorse technology for measuring both xenobiotic molecules and their metabolites in human tissues, as well as the body’s response to these molecules (e.g. via proteomics and metabolomics).^5, 15, 16^ MS measures ionized molecules in the gas phase as the ratio of their mass (m) to the number of charges (z) they carry (i.e. *m/z*). Molecules are typically extracted from samples prior to MS analysis, and the extracts are analyzed either directly or after an initial separation by gas or liquid chromatography or ion mobility spectrometry (IMS). Despite its exquisite sensitivity, resolution, and ability to measure thousands of peaks corresponding to unique chemical entities in exposomics and metabolomics studies, MS does not directly measure chemical structure. Instead, chemical structures are inferred through comparisons of experimental data to reference data libraries that have been generated through the analysis of authentic chemical standards.^17, 18^ However, the relevant chemical space of endogenous, anthropogenic, and other xenobiotic chemicals far exceeds the number of chemical standards available to the community.^5^ Further, the most commonly used property for MS-based molecular identification – the fragmentation (MS^2^) spectrum – is often inadequate for providing unambiguous identification of detected chemicals.^18-20^

Here, we evaluated if multi-dimensional spectrometric measurements coupled with comprehensive computational predictions of molecular properties could yield high confidence compound identifications without the use of authentic reference chemicals and within an extensive chemical space – so-called ‘reference-free’ compound identification. In this approach, several physicochemical properties are measured for a compound of interest, including accurate mass, tandem mass spectra, collision cross section, and infrared spectra; this ensemble of experimentally measured properties is then matched to computationally predicted properties. We selected fentanyl as a representative chemical of concern and demonstrated the feasibility of this methodology in a blinded analysis of a chemical mixture representative of a fentanyl tablet and against a chemical space comprising over 60 thousand known and chemically similar molecules and >1 billion computationally predicted analogs. The tablet contained 12 fentanyl analogs, 10 of which were readily flagged based on their observed properties. The 2 unflagged analogs were either not fully resolved or lacked the signatures used for flagging. Of the 10 flagged compounds, 6 were correctly identified using this methodology (i.e. the combination of measured properties matched best to the true positive analog from the 1+ billion candidates). One analog was identified as the second most probable candidate, while the remaining 3 were identified as the 5^th^, 7^th^, and 12^th^ most probable candidates. Overall, of the 10 suspected fentanyl features, 8 were correctly identified within the top 5 possible candidates from a starting list of >1 billion analogs. In post analysis, we find that all candidates could be identified within the top 5 plausible matches. This study demonstrates the feasibility of multi-dimensional spectrometric measurements and computational predictions of molecular properties for reference-free compound identification in exposome and related studies.

## Results

### Fentanyl as a representative chemical class with a broad chemical space

Fentanyl (*N*-phenyl-*N*-[1-(2-phenylethyl)piperidin-4-yl]propenamide) is a synthetic opioid first synthesized in 1959 and used as an anesthetic and analgesic.^21^ It is comprised of a central piperidine ring with an phenethyl group on the piperidine nitrogen and an N-phenylpropanamide at the piperidine 4-position. As an analgesic, fentanyl is 50-100 times more potent than morphine. Over 100 fentanyl analogs with varying potencies have been reported to date. For example, carfentanil differs from fentanyl by substitution of the 4-piperidinyl hydrogen with a carboxymethyl group and is 10,000 times more potent than morphine. Illicit use of illegally made fentanyls and other synthetic opioids has fueled the third wave of opioid overdose deaths in the U.S., with 82,000 deaths reported in 2022.^22^ Further, many users of illicit drugs are unaware that they are consuming fentanyl, as fentanyl is increasingly added to other substances or included in counterfeit pills.^21^ We chose to evaluate the reference-free compound identification concept using fentanyls as a representative chemical class of concern with extensive chemical diversity and began with computationally predicting the full, relevant chemical space (Figure 1).

**Figure 1.**
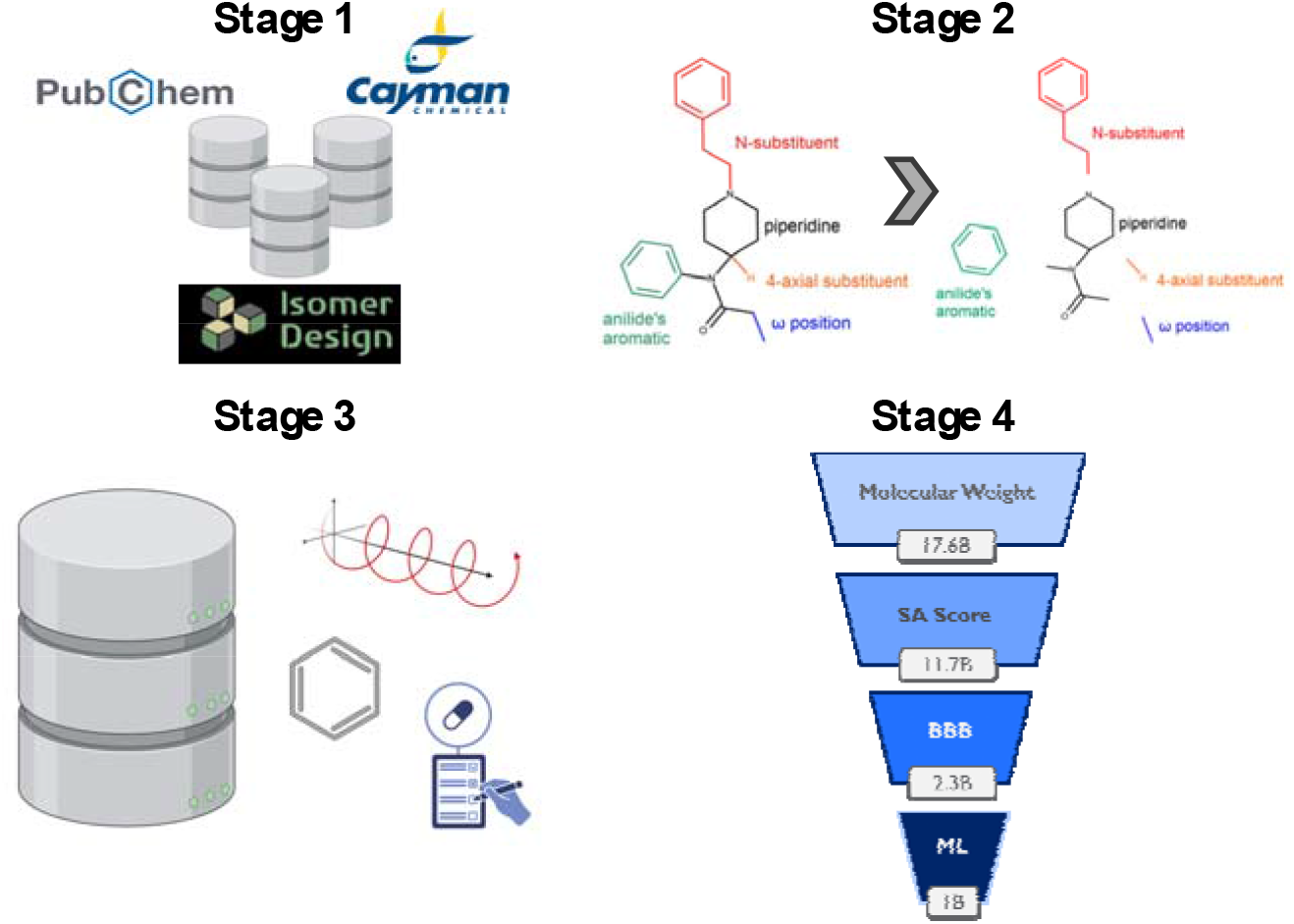
Computational workflow for predicting fentanyl analog chemical space. To create a comprehensive search space of all possible fentanyl analogs, structural information for existing fentanyl analogs were first downloaded from publicly available sources, including PubChem, Cayman Chemical, and Isomer Design (Stage 1). The structures of these existing analogs were computationally disassembled to substructures, and the substructures combinatorically recombined to yield over 1 trillion analog structures (Stage 2). Over 200 chemical properties (e.g. molecular polarizability, number of benzene rings, and quantitative estimate of drug likeness) were analyzed to inform a down selection strategy (Stage 3). The 1 trillion analogs were then down selected to just over 1 billion, based on exact mass, synthetic accessibility score, predicted blood-brain barrier permeability, and a consensus-based machine learning (ML) model trained on additional calculated chemical properties (Stage 4).

We began by compiling 61,451 structures of known fentanyls and chemically similar compounds, deconstructed these and enumerated their functional groups, and then combinatorically recombined these into a set of several billion computationally predicted analogs. These were then filtered based on exact mass, synthetic accessibility score, and predicted blood-brain barrier permeability, followed by final filtering using a consensus-based machine learning model trained on over 200 chemical properties, resulting in a filtered set of 1,001,957,837 novel, computationally predicted analogs. To create the final fentanyl chemical space, we combined the computationally predicted fentanyls with the 61,451 known fentanyl and chemically similar structures for a total of 1,002,019,288 unique fentanyl analog structures (we do not provide the structural information for the computationally predicted fentanyl analogs to avoid potential dual use^23^). We assessed the chemical similarity of the >1 billion fentanyl chemical space to fentanyl itself using the Tanimoto index^24^ analysis and found it to be remarkably high at 0.59, emphasizing the density of similar chemical structures represented by these analogs. By comparison, the average chemical similarity of fentanyl to 240,040 molecules within the Human Metabolome Database (HMDB) was 0.22, a chemical space with markedly lower density. Mass distributions for the fentanyl and HMDB chemical spaces are shown in Supplementary Figure S1.

### Multi-Dimensional Spectrometric Analysis of Mock Fentanyl Tablet

We next created a chemical mixture that simulated the composition of a fentanyl tablet. The mixture contained fentanyls, a chemically similar non-opioid molecule as a decoy (W-15), and three compounds that are typically used as cutting agents or in formulations of fentanyl tablets (mannitol, caffeine, and acetaminophen) at a 1000:1 ratio relative to the fentanyls. We added 12 different fentanyls to the mixture to challenge the combined resolution of the multi-dimensional spectrometric analysis and the corresponding accuracies of the computational predictions. Prior to analysis, seven tetralkylammonium salts (C2-C8) were added to the mixture as calibration compounds for mass and ion mobility measurements. The mock fentanyl tablet was then analyzed using an Orbitrap or Q-ToF mass spectrometer, both of which were retrofitted with structures for lossless ion manipulations (SLIM) devices.^25, 26^ The Q-ToF was further modified to allow for messenger tagging infrared spectroscopy after ion mobility separation (SLIM-cryoIR-ToF).^26^ The combination of measurements from the two instruments provided exact mass, MS^2^ spectra, collisional cross section (CCS), and infrared (IR) spectra for ionized molecules in the mixture, served to resolve between highly similar compounds, and concomitantly provided an ensemble of molecular descriptors. The mass and CCS domains were used to integrate data between the two instruments. During the study, members of the team who processed the data to identify the target fentanyls were kept blind to the composition of the mock fentanyl tablet. Thus, the results of experimental measurements and data processing that follow are reported from the perspective of a blinded analyst.

Figure 2 shows the results of the suite of measurements when applied to the mock fentanyl tablet. Figures 2a-b show the full mass spectrum as measured on the SLIM-Orbitrap and SLIM-cryoIR-ToF, respectively. A set of flagging criteria were used to identify *m/z* values that were suspected to be fentanyl analogs. Flagging criteria required the *m/z* values to display characteristics consistent with previous IMS-MS measurements of fentanyl^27-30^ and nitazene^31^ analogs, i.e. two CCS distributions whose respective MS^2^ spectra share some common fragment ions but otherwise are largely distinct. Using these criteria, a total of 10 *m/z* values (337.2278, 349.2278, 351.2431, 365.2587, 369.2337, 375.2068, 385.2040, 391.2743, 405.2906, and 429.2173) were flagged. Figures 2c-2f show the full set of measurements for the suspect *m/*z 375.2068 (subsequently referred to as ‘*m/z* 375’). This *m/z* is plotted in red in Figures 2a-b. Figures 2c-d show the extracted CCS distribution as measured on the two instruments. Supplementary Table S1 provides the *m/z*, CCS values, and the respective differences in measured values between instruments for the 10-suspect *m/z* targets. Calibration results in the *m/z* domain for both platforms are provided in Supplementary Table S2. Figures 2e-f display the MS^2^ and IR spectra measured for both CCS distributions, respectively. Previous studies suggest the two CCS distributions commonly observed for fentanyl analogs result from protonation site isomers.^32, 33^ In our post-analysis we explicitly confirm this, however, during the blinded study we exclusively used the IR spectrum of the larger CCS distribution for compound elucidation as its IR spectrum could be recorded at higher signal-to-noise ratios for all analogs.

**Figure 2.**
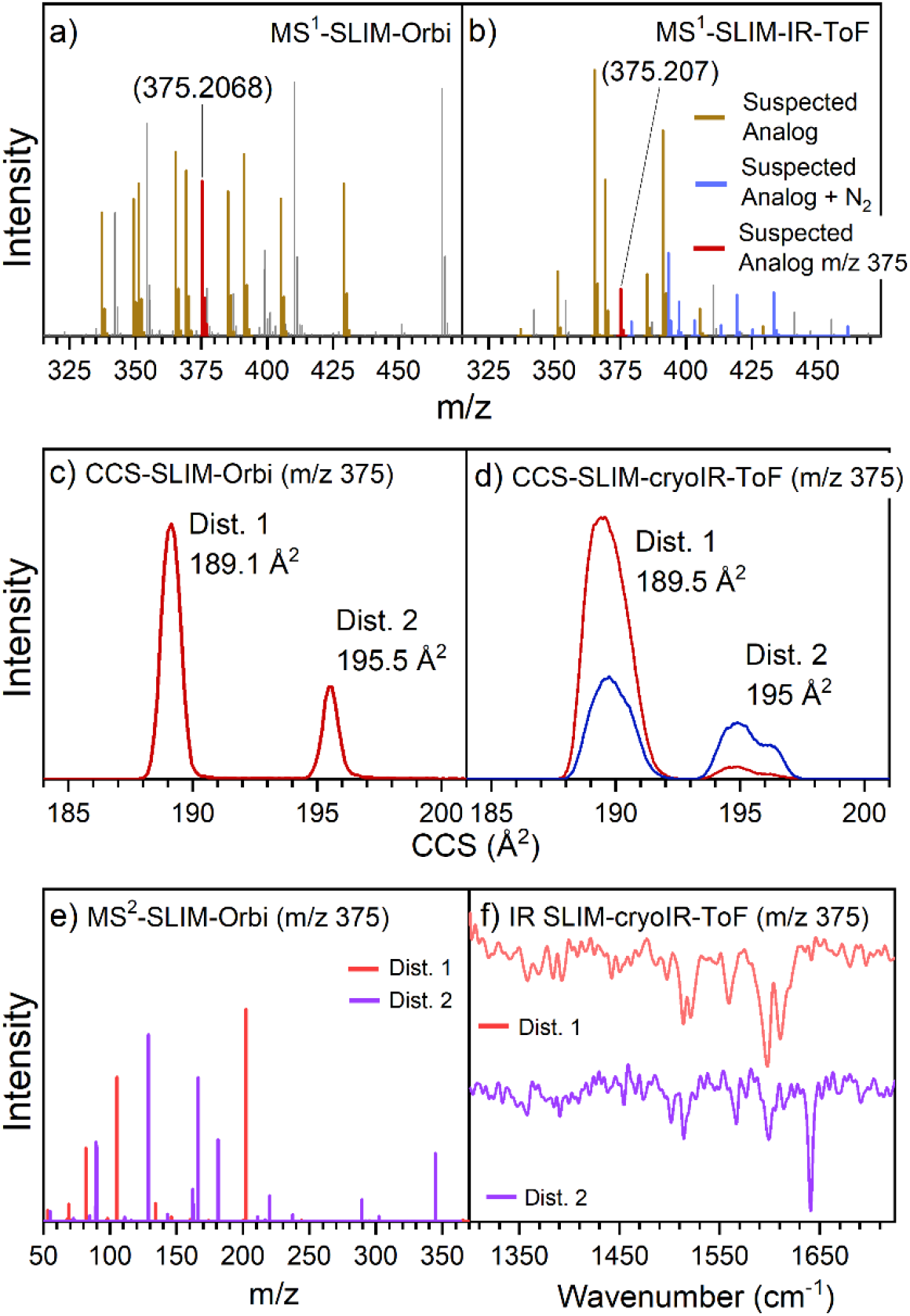
Ensemble of measurements made on the mock fentanyl tablet. MS measurements recorded on the SLIM-Orbitrap (HRMS) (a) and SLIM-cryoIR-ToF (b). Gray represents the background and calibrant ions, gold represents the m/z values that were flagged as fentanyl analogs, and blue are the m/z values of the N_2_-tagged, flagged fentanyl analogs. CCS values as measured on the SLIM-Orbitrap (c) and the SLIM-cryoIR-ToF (d) of the flagged nominal m/z 375. The blue trace in (d) shows the CCS distribution of corresponding N_2_-tagged population of nominal m/z 375. Tagging is observed selectively for the larger CCS distribution. MS^2^ spectrum recorded for both CCS distributions (e). IR spectrum recorded for the larger CCS distribution (f).

### Reference-Free Identification of Fentanyl Analogs

Figure 3 illustrates our reference-free compound identification paradigm. Molecular properties, which include CCS, MS^2^ and IR spectra, and exact mass measurements, are measured using hybrid mass spectrometers, as shown in Figure 2. Identical properties are then predicted from two- or three-dimensional chemical structures using machine learning (ML), quantum chemistry (QC), or molecular dynamics (MD). Compound identification is then accomplished by scoring the level of agreement between the measured and the predicted properties for all compounds of interest.

**Figure 3.**
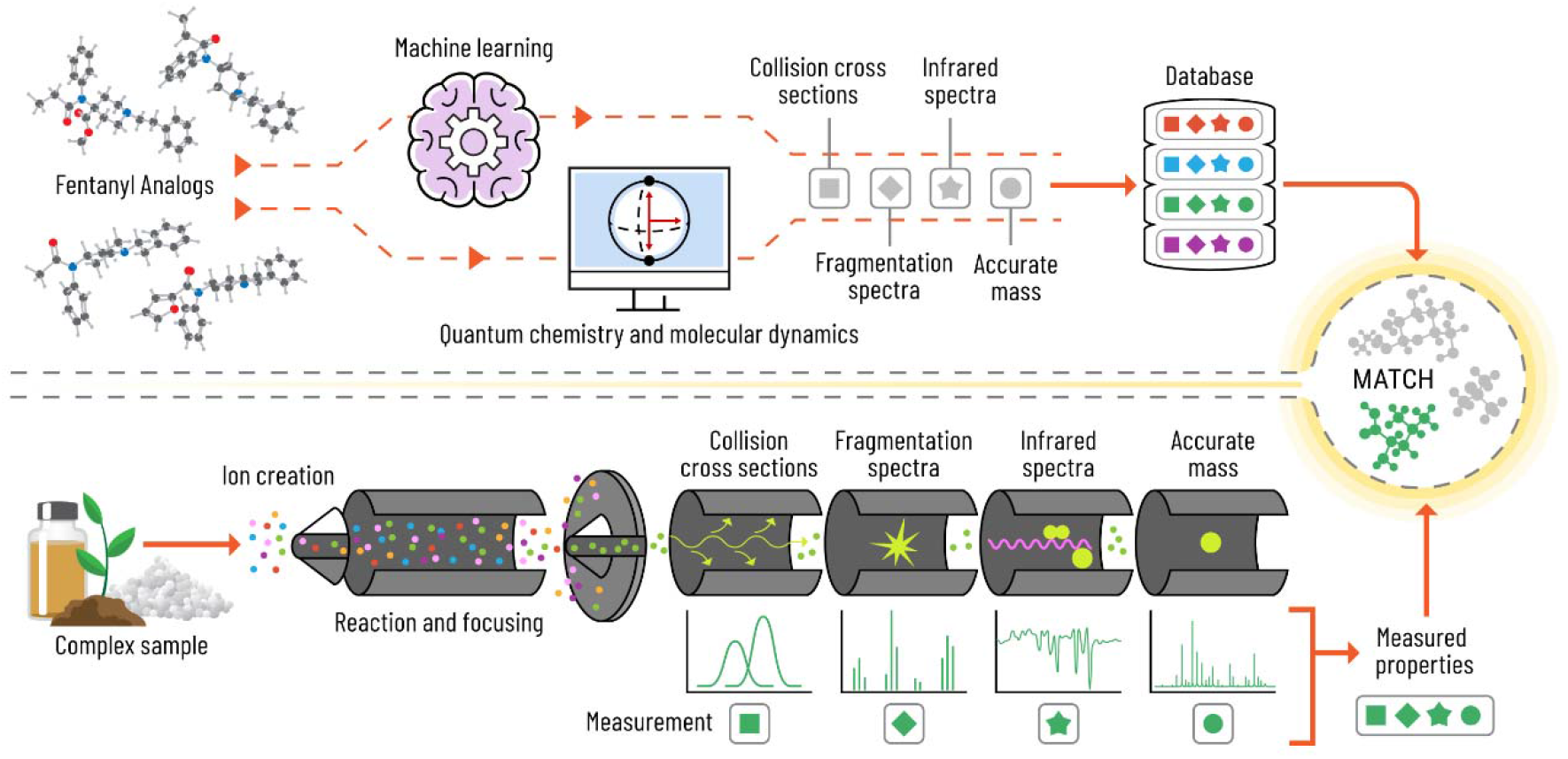
Reference-free workflow adopted in this work. The bottom panel shows ionization of a complex sample type into a mass spectrometer followed by several orthogonal measurements including CCS, tandem MS, IR spectra, and accurate mass for an ion of interest. Measured properties are stored in a database for future reference. The top panel shows the analogous predictive properties made for plausible input molecular structures. Computed properties are made using machine learning or first principles approaches and are stored into a computational reference library. Matching between experimental and computational observables allows for any measured compound to be identified. Adapted from Sarkar et al, Trends Mol Med, 2024 Dec;30(12):1137-1151 with permission from Elsevier.

Each molecular property provides information about a compound’s identity and can be computationally computed with varied levels of accuracy and precision. High resolution mass spectrometry measurements (Δ*m/z* < ∼2 ppm) allow potential candidates to be filtered or matched based on the agreement between their calculated (based on molecular formula) and experimentally observed masses. The ion mobility measurement, serves multiple purposes: it limits spectral overlaps in the mass domain, allowing for more accurate mass measurements; resolves between isomeric species that would otherwise be indistinguishable; and provides CCS values which can be calculated/predicted with an accuracy of > 97%.^34^ The MS^2^ domain provides a spectral signature that is representative of how an ion dissociates upon collisional activation. Fragment ions are useful for inferring how the compound is connected, provides information about chemical moieties that might be present, and can distinguish between isomers in favorable cases. The prediction accuracy of MS^2^ spectra varies across chemical classes. Lastly, the cryogenic IR spectrum serves as an additional precise spectral descriptor that provides information about the presence of certain functional groups, is highly sensitive to isomeric differences between compounds, and can be calculated with a high degree of accuracy in favorable cases. Our molecular property prediction pipeline is modular in nature, allowing for the exchange of algorithms and tools as their prediction accuracies are continually improved. In this work, MS^2^ spectra were predicted using QC-GN^2^oMS^2^,^35^ and CCS values were predicted in two stages: the first stage used a ML approach (sigmaCCS^34^) and the second stage used a MD approach (HP-Ω^36^). IR spectra were predicted from harmonic vibrational frequencies calculated by density functional theory (DFT) using ORCA 5.0.^37^

Reference-free identifications for all flagged *m/z* values were made in a tiered approach where analogs were scored and down selected by varied matched properties at each filtering stage until a single analog was selected as the most probable match. An overview of this process is provided in Figure 4 and was developed during the blinded analysis. The analysis of scores and implementation of thresholds was performed by manual analysis. In a post-analysis after unblinding, we performed a reanalysis to explicitly test this methodology’s accuracy without the bias introduced via blinded down selection.

**Figure 4.**
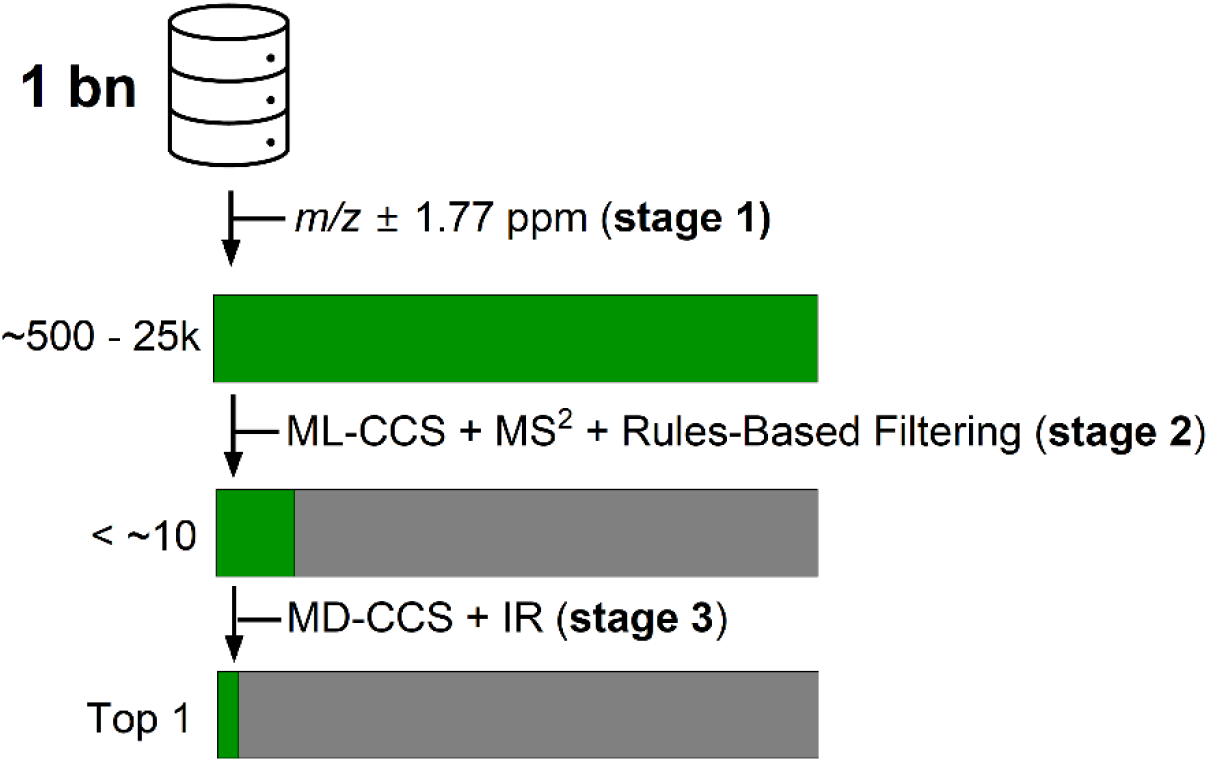
Overview of the down selection process employed in this work. First the suspect *m/z* is matched to the expected *m/z* of all candidates (1 bn) in the computationally generated library. Next, the ML-CCS and MS^2^ predicted values and spectra for mass-matched candidates are scored against experimentally measured values and filtered using a rules-based MS^2^ analysis. The top 10% of remaining candidates are then promoted to the final stage of matching to MD and DFT predictions. Here each candidate is represented with a max of 100 conformers. The calculated IR and CCS values for each conformer are then compared to the experimentally measured values. The isomer with the highest scoring conformer is selected as the most probable candidate. Green represents the distribution of plausible candidates while grey represents candidates that are filtered out after each down-selection step. The approximate number of remaining candidates after each down-selection step is provided at each filtering stage.

The blinded down selection process consisted of 3 main filtering stages as shown in Figure 4. Implementation of these stages are detailed in Figure 5, using the elucidation of suspect *m/z* 375 as an example. While this analysis was done in a blinded fashion, we highlight the scores of the true positive (TP) structure for clarity. First (stage 1), the measured mass of the flagged *m/z* was queried against the calculated masses of the ∼1 billion analogs present in the suspect list (Figure 5a). Analogs that matched the target *m/z* within ±1.77 ppm were retained. In the case of *m/z* 375, the suspect list was reduced to 722 candidates.

**Figure 5.**
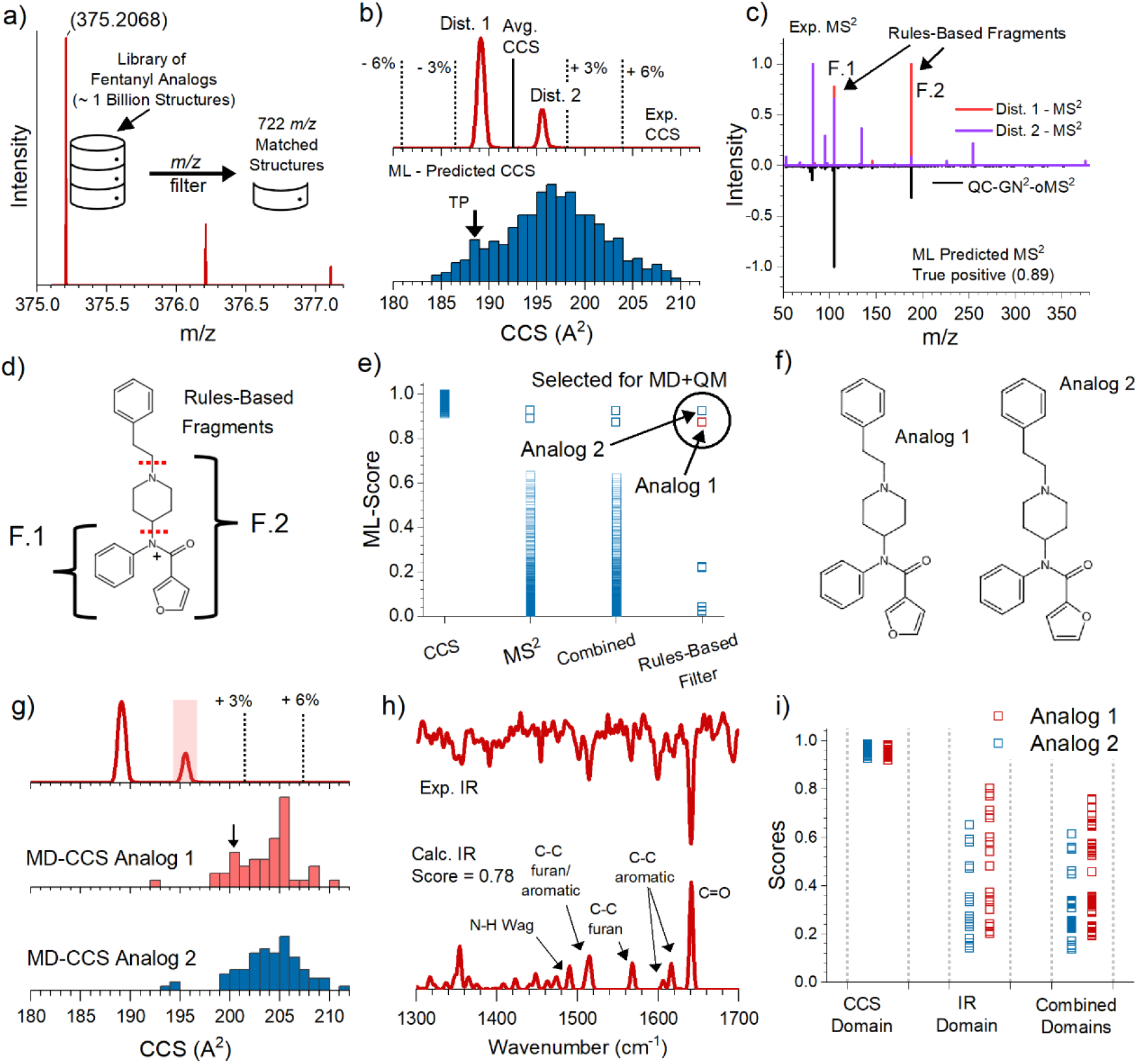
Down selection and identification process for flagged m/z 375. (a) HRMS measurement and mass matching against target library to reduce potential candidates from > 1bn to 722. (b) Experimentally measured CCS distributions (top) and histogram of ML calculated CCS values for all mass matched analogs (bottom). (c) Comparison of experimentally measured MS^2^ spectra for both CCS distributions (top) and the ML predicted MS^2^ spectrum of the true positive analog (bottom). (d) Cleavage sites for rules-based fragmentation analysis. (e) Summary of 722 ML scores in the CCS and MS^2^ domains, the combined scores, as well as the scores for the remaining candidates after the rules-based analysis. (f) Structures of the two highest scoring analogs (true positive and true negative) that are promoted to MD and QM modeling. (g) Comparison of experimentally measured CCS distributions (top) to the MD calculated CCS distributions for the ∼100 representative geometries of true positive (middle) and true negative analog (bottom). (h) Comparison of the experimental IR spectrum of the larger CCS distribution to the calculated IR spectrum of the highest scoring 3-D geometry of the ∼200 modeled geometries. (i) Summary of MD and QM scores in the CCS and IR domains of the representative geometries for the true positive and true negative analogs. The combined scores show the geometries of the true positive out score the geometries of the true negative allowing the identity of m/z 375 to be correctly elucidated.

In stage 2, the measured CCS and MS^2^ spectra of *m/*z 375 were scored against the ML-predicted values of the 722 remaining candidates. An additional MS^2^ rules-based analysis is also implemented at this stage to incorporate domain level knowledge. Figure 5b shows the comparison of the experimental (top) and ML-predicted (bottom) values for *m/z* 375 in the CCS domain. ML-predicted values are scored against the average CCS and composite MS^2^ spectrum of the two distributions, as we do not anticipate ML models to have the precision required to predict the difference between protomers. The experimental average CCS is indicated in the top panel of Figure 5b with a solid black line; dashed black lines indicate CCS values within ± 3 and ± 6 % of this value. ML-calculated values (Figure 5b bottom) for the 722 remaining analogs are displayed using a histogram and show that most of the predicted values fall within -3 to +6% of the average CCS value. In the CCS domain, each analog is scored using Equation 1:

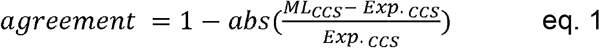

The black arrow in the bottom panel of Figure 5b indicates the bin that contains the TP analog of *m/z* 375, which has an agreement value of >0.97. The distribution of scores in the CCS domain is plotted in the first column of Figure 5e.

The MS^2^ spectra for both CCS distributions are plotted in the top panel of Figure 5c. ML predicted MS^2^ spectra are compared to the experimental composite MS^2^ spectra using a cosine similarity metric. The ML predicted spectrum of the TP of *m/z* 375 (score of 0.89) is shown in the bottom panel of Figure 5c. The distribution of cosine similarity scores in the MS^2^ domain is plotted in the second column of Figure 5e.

As we are working in the targeted chemical space of fentanyl, we evoke an additional metric in the MS^2^ domain for enhanced down selection. From the literature we can identify common fragmentation patterns in fentanyl and its associated analogs where cleavage commonly occurs at the amide bonds.^27^ These cleavage sites are highlighted in Figure 5d with red dashed lines and are referred to as rules-based fragments. Using the 2-D structure of each candidate analog, we calculated the *m/z* of the rules-based fragment ions (F.1 and F.2) formed when the indicated bonds are cleaved. The *m/z* values vary because of isomeric differences between candidate analogs. The number of *m/z* values in the experimental spectrum that match the *m/z* values of F.1 and F.2 were logged for each analog and appended as an additional score (max of 2). For reference, the TP analog of *m/z* 375 forms rules-based fragment ions with *m/z* values (nominal mass) of 105 (F.1) and 188 (F.2). The experimental MS^2^ spectrum, Figure 5c (top), shows two product ions that match these values, providing the TP analog with a rules-based score of 2.

While this filter will not always be possible to implement, it should be integrated when possible, to aid in selectivity and to alleviate the need for highly accurate ML-based property prediction models when they are not available.

At this stage of filtering, the TP analog of *m/z* 375 had ML-based property match scores in the CCS and MS^2^ domains of 0.98 and 0.89, respectively. These scores were multiplied to obtain a combined ML score of 0.87. This analog was further appended with a value of 2 to indicate the *m/z* values of its expected product ions from a rules-based fragmentation analysis of the MS^2^ spectrum.

The scoring summary for all 722 candidates for *m/z* 375 is plotted in Figure 5e. The scores in the ML-CCS domain were relatively high and provided little selectivity for resolving analogs. The scores in the MS^2^ domain on the other hand spanned a range of 0.00 to 0.92 and demonstrated an increased ability to resolve between potential matches. The third column shows the combined ML scores obtained after multiplying the scores from both domains. Because the CCS scores were all close to 1, the distribution of the combined scores strongly resembled the distribution of scores in the MS^2^ domain. The last column shows all analogs (9) that had scores of 2 in the rules-based analysis (end of stage 2). Analogs with a combined score within 10% of the highest scoring analog were promoted to stage 3; in the case of *m/z* 375, that included only the top 2 structures, circled in the last column of Figure 5e. The identities of the two promoted analogs are provided in Figure 5f, one of which is the TP while the other is a high scoring true negative (TN). The difference between the two top candidates is only in the position of the oxygen in the furan ring (position 3 vs. 2). At the end of stage 2, the TP and TN have similar scores, with the TN scoring slightly better.

The final filtering stage (stage 3) includes more rigorous MD- and DFT-based modeling of the selected analogs to predict their CCS and IR spectrum, respectively. The computational approach for this stage is detailed in the Methods section. In brief, each promoted analog is represented using a maximum of 100 conformers, each described by a calculated CCS value and IR spectrum that is matched to the experimental values of the flagged *m/z*. The agreement in the CCS and IR domains was evaluated using Equation 1 and a cosine similarity metric, respectively. A combined representative score was obtained by multiplying the score from both domains. The candidate that contains the highest scoring geometry was designated as the most probable match. Since we are readily able to record the IR spectrum of the second distribution, the calculated values for the CCS and IR spectrum are compared to the experimental values associated with the second CCS distribution (Dist. 2; highlighted in Figure 5g). Details regarding modeling of the protonation configuration during the blinded/unblinded analysis are provided in the Supplementary information (Supplemental Figures S2-S7).

Figure 5g shows the results of the MD-based CCS modeling for the two candidate analogs of *m/z* 375. The top panel reproduces the experimental CCS measurement for *m/z* 375. Dotted lines mark CCS differences of +3 and +6 % of this value. A histogram of the distribution of CCS values adopted by geometries of analogs 1 and 2 are shown in the middle and bottom panels of Figure 5f, respectively. Modeled CCS distributions are similar to one another, as can be expected based on the small difference between the two analogs. In both cases most of the modeled values fall below a 6% difference relative to the CCS value of the second distribution. The first two columns in Figure 5i show the MD-CCS scores for the conformers of both analogs. Figure 5h plots the experimentally recorded IR spectrum of *m/z* 375 (top) versus the calculated IR spectrum of the highest scoring (combined score) conformer (bottom; score of 0.78). The major normal modes that match vibrational bands in the experimental spectrum are labeled in the calculated spectrum. The second 2 columns in Figure 5i show the distribution of the scores in the IR domain of the representative conformers for both analogs.

### NOMINAL MASS OF SUSPECTED FENTANYL ANALOGS

Figure 5i shows the scoring summary for all conformers of the two candidate analogs. Like the ML-CCS scores, the MD-CCS domain showed an inability to resolve between the two analogs as all conformers scored better than 0.9. The IR scores showed increased conformer selectivity as the scores ranged between 0.14 and 0.80. More importantly, the IR domain shows the ability to resolve between analogs as the highest scoring conformer of analog 1 and 2 were 0.80 and 0.65, respectively. Since the CCS scores were all near one, the distribution of combined scores resembled the score distribution of the IR domain. We see several geometries adopted by analog 1 score higher than the highest scoring conformer of analog 2. The maximum combined score for both analogs was 0.61 (analog 2) and 0.76 (analog 1). It is important to note that the distribution of the combined scores is less important than the maximum score when the analysis is performed at the conformer level, as is done in this step. This is because the conformational sampling is designed to find multiple low-energy minima on the potential energy surface, rather than consistently exploring the lowest energy minimum. As a result, the conformers subjected for analysis ideally are representative of a large portion of the molecule’s potential energy surface but may be a sparse representation of a given minimum. As a result, we are less interested in the distribution of scores but rather the maximum score. From this full analysis, we assign the identity *m/z* 375 as analog 1 (3-furancarboxamide fentanyl), which was found to be the correct identity after unblinding.

The mock tablet contained 12 fentanyl analogs, and 10 were correctly flagged for elucidation via our workflow. The down selection analysis applied to the 10 flagged *m/z* values is provided in Supplemental Figures S8-S16. The identities, nominal *m/z* values, and 2-D structures of the 12 analogs used are provided in Supplementary Figure S17. The two analogs not flagged were remifentanil (m/z 377.2075) and phenyl fentanyl (m/z 385.2280), see Supplementary Information, Figures S18-S21.

The nominal masses and identification results for the 10 flagged analogs are provided in Table 1, along with the number of potential analog matches remaining after each down-selection step. Mass filtering (stage 1), as can be expected, provided the largest reduction (1+ billion down to several thousand) in potential candidates. Candidates remaining after this stage consisted mostly of isomers. ML predictions of MS^2^ spectra along with rules-based fragmentation analysis (stage 2) provided the next largest reduction in potential candidates, several thousand down to double digits or less. Finally, the MD CCS and IR analysis (stage 3) provided orthogonal information to select between the remaining candidates, allowing for the best match to be selected. While there was not an explicit down selection in this stage, we do show the numbers of candidates remaining with IR match scores above 0.65 to provide insight into the selectivity afforded by the IR domain. A threshold of 0.65 was chosen based on comparisons between spectra to known standards from existing literature and previous experiments. After unblinding, we observed that the average IR score of the TP was 0.72.

**Table 1.** Summary of blinded and unblinded analysis. Nominal masses for all flagged m/z values are provided along with the number of remaining candidates after each down selection step. Penultimate row and last row indicate the rank of the TP after blinded and unblinded analysis, respectively. (T-# indicates the number of tied candidates in terms of rank score)

NOMINAL MASS OF SUSPECTED FENTANYL ANALOGS
| # OF PLAUSIBLE CANDIDATES REMAINING | 337 | 349 | 351 | 365 | 369 | 375 | 385 | 391 | 405 | 429 |
| --- | --- | --- | --- | --- | --- | --- | --- | --- | --- | --- |
| MASS FILTER (STAGE 1) | 806 | 813 | 2,222 | 5,187 | 3,287 | 722 | 22,233 | 13,235 | 22,255 | 7,007 |
| ML SCORE + DIRECTED FRAGMENTATION SEARCH (STAGE 2) | 4 | 5 | 28 | 26 | 6 | 2 | 3 | 3 | 1 | 1 |
| MD CCS + IR MATCH SCORE ABOVE 65% (STAGE 3) | - | - | 13 | 10 | 5 | 1 | 2 | 2 | 1 | - |
| TRUE POSITIVE ID RANK (BLINDED) | 1 | 5 | 12 | 7 | 2 | 1 | 1 | 1 | 1 | 1 |
| POST-ANALYSIS (UNBLINDED) | 1 | 5 | 1(T-6) | 4(T-3) | 2 | 1 | 1 | 2 | 1 | 1 |

The penultimate row in Table 1 indicates the rank of the TP analog as determined by our down-selection methodology while blinded. A rank of 1 indicates the correct analog was identified as the highest scoring and therefore most plausible analog among the potential candidates. IR spectra for the flagged *m/z* values of 337, 349 and 429 could not be recorded due to unoptimized transmission through the SLIM-cryoIR-ToF instrument. In those cases, identifications were made after the ML and rules-based analysis (Figure 4, filtering stage 2).

The last row shows the ranked scores of the 10 flagged analogs when reanalyzed after unblinding. Reanalysis was done using a data-driven approach to explicitly test the accuracy of our approach without bias from the blinded down selection process. We first established where the TP scored in terms of percentage relative to the top scores within the ML-CCS, MS^2^, and IR domains. The TP scores were found to be, at a minimum, within the top 97.9%, 93.4%, and 94.7% for the ML-CCS, MS^2^, and IR domains, respectively across analogs. During this reanalysis we excluded the MD-CCS domain as errors were found to be > 3%. For each domain we rounded the minimum percentage down to the nearest 0.5% as the cut-off criterion in each domain. Re-analysis was done reassessing the pool of plausible analogs at the end of stage 2. These candidates were filtered based on the data-driven cut-offs. The top candidate was then selected based on the highest score in the IR domain. The result of this re-analysis is provided in the last row of Table 1 and shows that all TP’s fall within the top 5 plausible candidates. All scores and filtering metrics are provided in a Supplementary Excel file.

## Discussion

In a traditional MS-based omics approach for molecular identification, the measured properties of detected molecules would be compared to reference values obtained through analysis of pure compounds. This means that only previously measured compounds can be identified, leaving novel molecules unidentified. In this study, we evaluated a reference-free approach for the identification of fentanyl analogs measured using a multi-dimensional spectrometric approach coupled with computationally predicted reference values. Additional orthogonal and semi-orthogonal measurements were used for resolving compounds with very similar chemistries. Because we were limited to using Drug Enforcement Administration-exempt preparations of fentanyls in our study, we did not expect any identifications to correspond to the >1 billion novel analogs that were computationally predicted and formed the basis of the suspect list. However, to fully evaluate and validate our reference-free identification approach, we thoroughly populated the relevant chemical space. In so doing, we effectively steered the analysis bias away from previously observed compounds, greatly expanded the considered chemical space, and inherently increased the number of possible false positives. It is important to note that the experimental observables measured in our study were strictly compared to computed properties. That is, no experimental spectra were compared to previously generated experimental spectra. We do however acknowledge that this is a targeted study that leveraged domain level expertise when analyzing the MS^2^ and CCS domains.

We have recently proposed a framework for evaluating compound identifications in untargeted metabolomics, which integrates observed molecular properties, measurement capabilities, and the complexity of the chemical space spanned by reference libraries supporting annotation to inform confidence in molecular annotations.^19^ Identification probability, the inverse of the number of candidate metabolite identifications for a given observed feature, is used to quantify identification confidence and is strongly dependent on measurement capabilities as well as the depth and breadth of chemical space represented within reference libraries.^38^ The present work introduces distinct challenges to applying this identification probability framework: the reference library is formed from near-exhaustive sampling of analogs of a single target compound, making it both far narrower in scope but deeper in coverage than typical reference libraries used for identification in untargeted metabolomics. The scale of candidate metabolite annotations for each measured feature would make the identification probability metric difficult to interpret, and the in-depth analyses of identifications that we had already performed were more informative, so we opted not to include identification probability in the present assessment.

The mock fentanyl tablet contained 12 fentanyl analogs mixed with a variety of cutting agents and a decoy compound. With our methodology, we were able to flag 10 analogs as likely fentanyls and were able to correctly identify 70% within the top 2 candidates. Importantly, identifications were performed using a suspect list of >1 billion possible analogs who’s average Tanimoto index score was 0.58. This level of accuracy is unprecedented for reference-free identification in such a congested chemical space and hinges on the multi-spectrometric measurements and corresponding ensemble of computational predictions used. The work presented here demonstrates the ability to precisely describe a compound using an ensemble of orthogonal and semi-orthogonal measurements coupled with computational predictions.

## Online Methods

### Prediction of fentanyl chemical space

The computational workflow for predicting all possible analogs of fentanyl consisted of 4 Stages (**Figure 1**): 1) compilation of existing fentanyl analogs and chemically similar compounds into an in-silico library, 2) enumeration and recombination of *in silico* library analogs to create new analogs, 3) structure and property-based analysis of analogs to inform filtering techniques, and 4) down selection of fentanyl analog chemical space

In Stage 1, we scraped open web sources and available libraries to build a large database of as many known and reported fentanyls and chemically similar compounds as possible, as previously described.^39^ This resulted in over 61,451 structures – represented in Simplified Molecular Input Line Entry System (SMILES)^40^ notation – that formed the library of fentanyl analogs and chemically similar structures used here as the reference compound list.

In Stage 2, all chemical substructure groups in the existing fentanyl analogs and chemically similar compounds were identified and a combinatorial analysis used to generate new analogs. Here, the analogs were converted to a SMILES Arbitrary Target Specification (SMARTS)^42^ representation, which provides a more straightforward method to identify chemical substructures. To disassemble the analogs into substructures, the fentanyl structural backbone and four substituent groups were first defined and the core saved as a SMARTS string. Next, the 61,451 fentanyl analogs from Stage 1 were decomposed into backbone and substituent groups, and unique substituent groups were enumerated at each of the four attachment points. This analysis identified 5,993 unique substructures at the N-substituent point, 404 unique substructures at the anilide aromatic point, 4536 substructures at the omega position point, and 56 substructures at the 4-axial substituent point. Approximately 700 billion new analogs were formed via recombination of substructures.

In Stage 3, the structures of the new analogs were converted to SMILES to computational facilitate property-based analysis to inform down selection techniques. While all subsequent analyses were performed on supercomputing resources, the computational time and resource requirements for a 700-billion-member dataset were still too high for most algorithms. As such, we analyzed the dataset to determine the least computationally expensive techniques to use for analysis and filtering steps. To assist in the process, we created a reference set of fentanyl analogs using the International Narcotics Control Board’s (INCB) set of fentanyl-related substances with no known legitimate uses.^43^ This set allowed us to establish reference ranges for over 200 calculated chemical properties (e.g., molecular weight, quantitative estimation of druglikeness (QED), topological polar surface area, number of H-bond donors and acceptors, etc.) within which the same properties for the new analogs were required to reside. A set of decoy compounds was also generated using the DUD-E^44^ web tool for use as a negative control. The combined datasets enabled the training of machine learning models to predict whether a given analog was chemically similar to fentanyl based on these calculated properties.

In Stage 4, we filtered the list of new analogs based on exact mass, synthetic accessibility score, and predicted blood-brain barrier permeability. Compounds remaining after these first three property-based filters were then assessed with a consensus-based machine learning (ML) model trained on additional calculated chemical properties. Filtering steps along with the approximate number of generated fentanyl analogs remaining following each filter are depicted in Figure 1. This down selection process ultimately reduced the set of 700 billion putative analogs to 1,001,957,837. A molecular docking simulation pipeline using AutoDock Vina^45^ was also implemented to interrogate binding potential at the m-Opioid Receptor (MOR). Our analysis of the pipeline’s ability to accurately predict binding affinity showed that while there is a statistically significant difference in binding affinity scores overall when comparing known dangerous fentanyls to decoys, individual scores are frequently unreliable. As such, we omitted the docking screen from the final filtering steps.

### Prediction of fentanyl molecular properties

Property predictions made for the candidate analogs included their exact mass, tandem mass spectra, CCS, and IR spectra. Throughout the text we refer to properties as ‘predicted’ regardless of the tool used (i.e. ML, MD, or DFT). While this is done for conciseness, we acknowledge that values obtained from ML tools are better classified as ‘predicted’ properties, while those obtained from MD and DFT tools are better classified as ‘calculated’ properties. Throughout the text we clarify which type of tool is used at each filtering stage but refer to all predicted/calculated properties as ‘predicted’ values. For all compounds, exact masses (Figure 4, Filtering stage 1) were calculated from their SMILES string using the open-source cheminformatics library RDKit (www.rdkit.org). ML CCS and MS^2^ values for all mass-matched analogs (Filtering stage 2, Figure 4) were calculated using sigmaCCS^34^ and QC-GN^2^oMS^2^, respectively. The version of QC-GN^2^oMS^2^ used in this work was specifically trained on fragmentation data from known fentanyls^35^. A custom Python script was implemented to calculate the masses of the rules-based fragment ions for all mass matched candidates based on SMARTS patterns describing common elements of fentanyl analog structures. Analogs that were promoted to filtering stage 3 (see Figure 4) were converted to a 3-D geometry from their 2-D SMILES strings using Open Babel.^46^ Protonation was performed using the protonation feature in Open Babel or through manual addition as needed. The CREST package^47^ was then used to perform a conformational search for each 3-D geometry using extended tight binding (xTB), generating between 50-200 conformations per input (input line provided in the Supplementary Information). The lowest energy structures (max 100) at the xTB level of calculation were then further optimized using density function theory (DFT) implemented via the ORCA package^37^. IR spectra were calculated for each conformer after geometry optimization (both performed at the B3LYP-D3(BJ)/def2-TZVP level of theory. The Orca input script is provided in Supplementary Information. The optimized geometries along with their Mullikan partial charges were used as inputs for the MD based CCS calculations performed using HP-Omega^36^.

### Matching predicted and experimental molecular properties

The masses of fentanyl analogs in the suspect chemical space were calculated based on the corresponding molecular formula and then compared to the masses of measured fentanyls within ±1.77 ppm based on the experimentally determined mass measurement accuracy. ML and MD CCS values were compared using eq. 1 where the experimental reference value was the average CCS or the CCS of the larger distribution, respectively. A cosine similarity metric was used to evaluate the level of agreement between the composite MS^2^ spectrum and each ML predicted MS^2^ spectrum. A cosine similarity metric was also used to evaluate the similarity between experimentally recorded and calculated IR spectra. All calculated IR spectra were scaled by 0.986 prior to being compared to experimentally observed spectra. The scaling factor was obtained empirically by subjecting the assigned [YGGFL+H]^+^ conformer from literature^48, 49^ (XYZ coordinates provided in the Supplementary Information) to the same level of DFT optimization used in this work. The resulting IR spectrum was then scaled such that the highest frequency stretch in the amide I region in the calculated IR spectrum (C=O stretch of the COOH group, unscaled frequency 1749 cm^-1^) was aligned to the corresponding highest frequency stretch in the experimental IR spectrum (1725 cm^-1^), Supplemental Figure S21, scale factor equals 0.986. The comparison of the experimental and scaled calculated IR spectrum of YGGFL is provided in the Supplementary Information.

IR spectra were only collected for distribution 2 in the CCS domain (Figure 2d and Figure 5b) due to preferential tagging (see Multi-dimensional spectrometric measurements section for details). To determine the most plausible protonation site associated with distribution 2, we performed a series of calculations. First, we assessed the final single point and Gibbs Free energy of ∼100 conformations of unsubstituted fentanyl in 4 protonation configurations when optimized in dielectrics equal to 100% water, 100% methanol (solvents used for analysis) and in the gas phase, Supplementary Figure S2. Calculations were performed using the Orca package, the relative energies in water and methanol were calculated using the Universal Solvation Model and Conductive Polarizable Continuum Models. Full thermodynamic calculations were performed for gas phase analysis and relative Gibbs Free energies were evaluated after IR frequencies were calculated. The Orca input scripts for these calculations are provided in Supplementary Information.

Further, we employed QCxMS^50^ (MD based MS^2^ predictions) and confirmed protomer 2 and 3 fragment differently from one another and largely accounts for the fragmentation differences observed experimentally between the two CCS distributions, (Supplementary Figure S4).

### Multi-dimensional spectrometric measurements

Drug Enforcement Agency-exempt preparations of fentanyl hydrochloride (CAS: 1443-54-5), β-methyl fentanyl hydrochloride (CAS: 1443-43-2), ortho-fluroisobutyryl fentanyl hydrochloride (CAS: 2748591-21-9), (±)-cis-3-methyl butyryl fentanyl (CAS: 88641-20-7), furanyl fentanyl (3-furancarboxamide isomer) hydrochloride (CAS: 2306823-47-0), para-chloroisobutyryl fentanyl hydrochloride (CAS: 2306827-85-8), cyclohexyl fentanyl hydrochloride (CAS: 2309383-14-8), 2,2,3,3-tetramethyl cyclopropyl fentanyl hydrochloride (CAS: 2309383-12-6), phenyl fentanyl hydrochloride (CAS: 2309383-16-0), remifentanil hydrochloride (CAS: 132539-07-2), cyclopropyl fentanyl hydrochloride (CAS: 2306825-44-3), benzodioxole fentanyl hydrochloride (CAS: 2306823-01-6), and W-15 (CAS: 93100-99-3) were purchased from Cayman Chemical (Ann Arbor, MI, USA) and prepared at 1-10 µM equimolar mixtures in 50/50 methanol/water. Seven tetralkylammonium salts (C2-C8) were prepared in the same sample at 100 nM equimolar concentration and served as calibration compounds for the mass and mobility measurements. Three additional compounds (mannitol, caffeine, and acetaminophen) that are typically used as cutting agents or in formulations of fentanyl tablets were also included in the mixture at 1 mM equimolar concentration to simulate a real-world analysis and provide potential false positive/miscellaneous signals. These cutting agents were purchased from Millipore-Sigma (St. Louis, MO, USA). The final fentanyl mixture contained a total of 24 compounds. The identity of the mixture’s constituents was kept from the individuals tasked with processing the data and making the final assignments.

All measurements were made using either a Q-Exactive Plus Orbitrap (Thermo-Fisher, Waltham, MA, USA) or Agilent 6538 Q-ToF mass spectrometer, both of which were retrofitted with structures for lossless ion manipulations (SLIM) devices to allow for high-resolution ion mobility spectrometry measurements before other (e.g. mass) analyses.^25, 26^ The Agilent 6538 Q-ToF was further modified to allow for messenger tagging infrared spectroscopy after the mobility separation.^26^ The combinations of these two instruments provided exact mass, tandem mass spectra, collisional cross section, and infrared spectra measurements. The full instrument descriptions can be found elsewhere.^25, 26^ Briefly, ions were generated using micro-flow electrospray ionization at flow rates of < 1 µL/min. Emitters were fabricated by dipping fused silica emitters into aqueous hydrofluoric acid as described elsewhere. After ionization, on both platforms, ions were collimated by dual funnels and guided into a 13 m SLIM device held at ∼ 2.5 torr of N_2_. The operation and principles of this technology are described in detail elsewhere. Experiments begin with a short in-SLIM accumulation where ions were accumulated in the device at low traveling wave amplitudes (< 5 V_pp_) for 50 -200 ms. After the accumulation, the traveling wave amplitudes were raised to separation conditions (20-30 V_pp_) to initiate the mobility separation. As ions transversed the SLIM device and responded to the oscillating electric fields, they separated by mobility and eventually exit the device as mobility-separated packets. In the SLIM-Orbitrap, mobility-separated packets were gated again (using a dual-gate) prior to mass and tandem mass analysis, as described previously.^25^ In the modified Agilent 6538 Q-ToF (SLIM-IR-MS), mobility separated ion packets transverse a cryogenically (30 K) held SLIM where they were collisionally cooled for messenger tagging spectroscopy. In this device, well-resolved infrared spectra were acquired for mobility separated and tagged compounds.^26^ The mass and mobility information were matched between instruments to obtain a full set of descriptors for a given compound of interest, which were then stored in a custom library for referencing.

## Data availability

The raw mass spectrometry (or experimental) data have been deposited to the MassIVE repository (massive.ucsd.edu) with the dataset identifier MSV000101535.

## Code availability

All code used for predicting molecular properties was described in prior publications and previously made available. Code to generate novel fentanyl analogs combinatorially will not be released due to dual-use concerns.

## Acknowledgments

We thank the Pacific Northwest National Laboratory (PNNL), Laboratory Directed Research and Development (LDRD) Program for supporting this research via the *m/q* Initiative. Multimodal experimental analyses were performed in the Environmental Molecular Sciences Laboratory, a national scientific user facility sponsored by the U.S. Department of Energy, Office of Biological and Environmental Research. We also thank the following individuals for their helpful guidance through the course of developing the computational and experimental capabilities and in their demonstration for the fentanyls use case: Dr. Peter Armentrout, University of Utah; Dr. Augustus Fountain, University of South Carolina; Ms. Rachel Gooding, Department of Homeland Security; Dr. Stefan Grimme, University of Bonn; Dr. Tobias Kind, Enveda; Dr. Thomas Rizzo, Swiss Federal Institute of Technology; Dr. Kabrena Rodda, PNNL; and Dr. Akos Vertes, The George Washington University. Lastly, we thank Mr. Nathan Johnson of PNNL for graphics assistance with figures. PNNL is operated by Battelle Memorial Institute for the U.S. Department of Energy under Contract DE-AC05-76RL01830.

## Funding

This research was supported by the PNNL Laboratory Directed Research and Development (LDRD) Program and is a contribution of the *m/q* Initiative (https://www.pnnl.gov/projects/mq-initiative). Additional support was provided by the Predictive Phenomics Initiative (https://www.pnnl.gov/projects/predictive-phenomics-science-technology-initiative) via the PNNL LDRD Program.

## Author Contributions

C.P.H – Conceptualization, data curation, formal analysis, investigation, methodology, resources, software, supervision, validation, visualization, writing-original draft, writing-review & editing

A.L.H - Conceptualization, data curation, investigation, methodology, resources, software, supervision, visualization, writing-original draft

D.C – Conceptualization, data curation, methodology, resources, software, supervision, validation, visualization, writing-review & editing

K.J.S – Conceptualization, data curation, formal analysis, investigation, methodology, resources, software, supervision, validation, visualization, writing-original draft, writing-review & editing

R.O – Conceptualization, methodology, data curation, methodology, resources, software, validation, writing-review & editing

P.S.R – Data curation, investigation, methodology, software, validation, writing-review & editing

E.K. – Data curation, software, validation

J.N - Data curation, software, validation

D.H.R- Conceptualization, methodology, visualization, writing-review & editing

V.S.L – Resources, writing-original draft, writing-review & editing

G.Y.D - Data curation, software, visualization

E.B. - Data curation, validation

B.M.W – Conceptualization, project administration

S.R – Conceptualization, methodology, supervision

Y.M.I – Conceptualization, methodology, supervision, writing-review & editing, project administration

R.G.E – Conceptualization, funding acquisition, supervision, visualization, project administration

T.O.M – Conceptualization, methodology, funding acquisition, project administration, supervision, visualization, writing-original draft, writing-review & editing

## Ethics Declaration

The authors declare no competing interests.

## Notes

### Competing Interest Statement

The authors have declared no competing interest.

