## Supplementary for "Reference-free compound identification using computational prediction of molecular properties and multi-dimensional spectrometric measurements: a fentanyl case study"

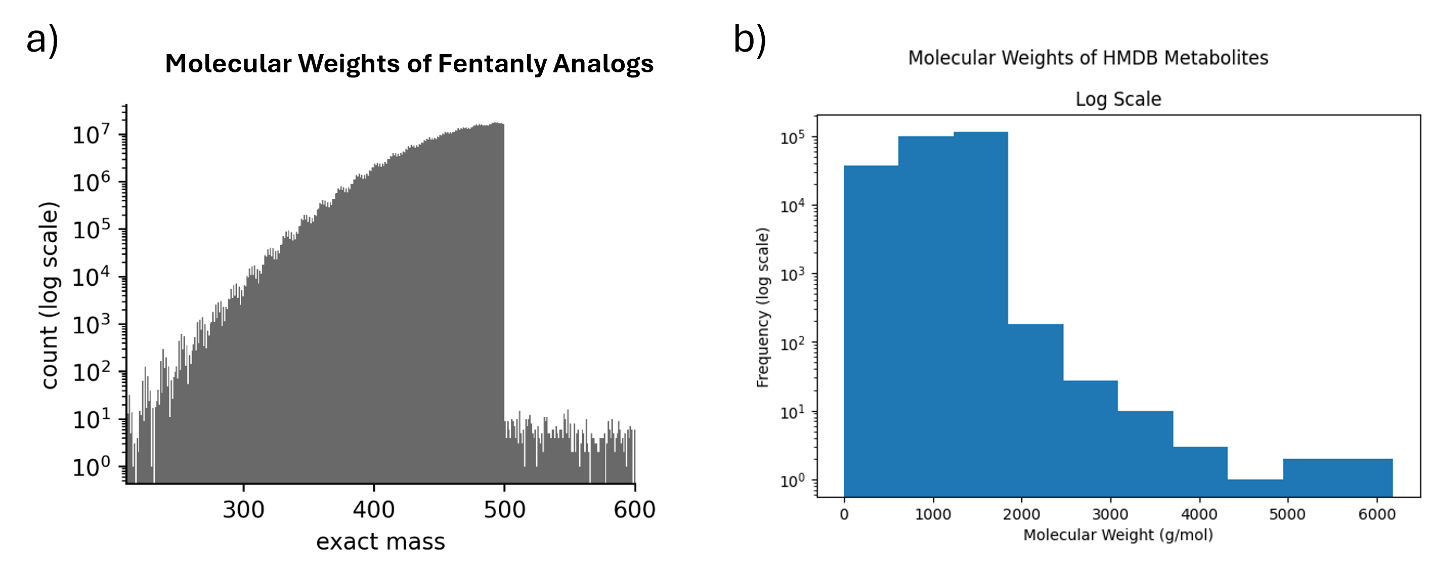

**Supplementary** **Figure S1.** Mass distributions for the fentanyl and Human Metabolome Database chemical space used for Tanimoto Index analysis. A) fentanyl chemical space, b) HMDB chemical space.

**Table S1. Measured m/z and CCS values from the SLIM-Orbitrap and SLIM-cryoIR-ToF for the 10-suspect m/z values and the relative agreement between measured values.** Two CCS values and associated differences are reported, corresponding to the two experimentally observed arrival time distributions for fentanyl analogs.

| SLIM-Orbitrap | | | SLIM-IR-ToF | | | Agreement | | |
| --- | --- | --- | --- | --- | --- | --- | --- | --- |
| m/z | **CCS (Å^2^)_1_** | **CCS (Å^2^)_2_** | **m/z** | **CCS (Å^2^)_1_** | **CCS (Å^2^)_2_** | **Δ m/z (ppm)** | **Δ CCS (%)_1_** | **Δ CCS (%)_2_** |
| 337.2272 | 181.9 | 187.5 | 337.225 | 183.9 | 187.6 | 6.52 | 1.10 | 0.05 |
| 349.2273 | 185.3 | 191.8 | 349.224 | 186.4 | 192 | 9.45 | 0.59 | 0.10 |
| 351.2432 | 185.42 | 192.3 | 351.239 | 186.4 | 192.5 | 11.96 | 0.53 | 0.10 |
| 365.2585 | 188.3 | 194.3 | 365.26 | 188.8 | 193.5 | 4.11 | 0.27 | -0.41 |
| 369.2339 | 186.6 | 193.7 | 369.228 | 187.1 | 192.9 | 15.98 | 0.27 | -0.41 |
| 375.2068 | 189.1 | 195.5 | 375.2074 | 189.4 | 195 | 1.60 | 0.16 | -0.26 |
| 385.2039 | 194.6 | 201.6 | 385.207 | 194.4 | 201.1 | 8.05 | -0.10 | -0.25 |
| 391.2745 | 197.4 | 205.7 | 391.273 | 197.2 | 204.9 | 3.83 | -0.10 | -0.39 |
| 405.2899 | 201.5 | 210.4 | 405.279 | 201.6 | 209.7 | 26.89 | 0.05 | -0.33 |
| 429.2177 | 203.5 | 211 | 429.209 | 203.36 | 211.1 | 20.27 | -0.07 | 0.05 |

**Table S2. m/z calibration results obtained for tetra-alkyl ammonium salts (TAA) for the Orbitrap and ToF mass analyzers**

| Orbitrap Mass Analyzer | | | | |
| --- | --- | --- | --- | --- |
| TAA Theoretical Mass | **TAA Measured Mass** | **Abs(Delta)** | **Mass Error (ppm)** | **Average Mass Error (ppm)** |
| 242.2846 | 242.2841 | 0.0005 | 2.0637 | 1.7705 |
| 298.3472 | 298.3467 | 0.0005 | 1.6759 |  |
| 354.4099 | 354.4093 | 0.0006 | 1.6930 |  |
| 410.4726 | 410.4719 | 0.0007 | 1.7054 |  |
| 466.5356 | 466.5348 | 0.0008 | 1.7148 |  |
| ToF Mass Analyzer | | | | |
| TAA Theoretical Mass | **TAA Measured Mass** | **Abs(Delta)** | **Mass Error (ppm)** | **Average Mass Error (ppm)** |
| 242.2846 | N/A | N/A | N/A | 7.2189 |
| 298.3472 | 298.3464 | 0.0008 | 2.6814 |  |
| 354.4099 | 354.4112 | 0.0013 | 3.6681 |  |
| 410.4726 | 410.4718 | 0.0008 | 1.9490 |  |
| 466.5356 | 466.526 | 0.0096 | 20.5772 |  |

**Protonation Site Analysis (blinded):**

Data from reference 27 and 33 in the main text show fentanyl and related analogs display two IMS distributions during IMS-MS analysis. Upon fragmentation, it is observed that the distributions fragment differently. This phenomenon is consistently observed across analogs. Further, it was observed that the ratio of the two distributions is sensitive to the solvent conditions used for electrospray, and the two distributions cannot be forced to interconvert in the gas phase when exposed to intentional ion activation within the SLIM device. These observations strongly suggest that the two distributions are protonation site isomers.

During the blinded analysis, specifically at stage 3 of the down selection process, we selectively use the observables of the larger CCS distribution for identification. This is done as the IR spectra consistently has higher single-to-noise ratios compared to the IR spectra collected for the smaller CCS distribution. As a result, we must explicitly define the protonation site associated with the larger CCS distribution to confidently compare the predicted values to the experimental observables. To elucidate the protonation configuration associated with the larger CCS distribution, we performed a series of computations on unsubstituted fentanyl (m/z 337). First, we investigated the plausible protonation sites and their relative energies and then looked at MD simulations of CCS and MS^2^ spectra of the most energetically accessible protomers. Finally, we analyzed the experimental IR spectral differences between the two distributions to further validate our choice in the protonation configuration. We did this analysis blinded to maintain the integrity of the initial work which was to determine if a series of fentanyl analogs could be structurally elucidated via our reference-free paradigm. In our post-analysis, we explicitly confirm that our choice in protonation configuration is indeed correct by comparing the IR spectra of the true positives to the IR spectra collected for both CCS distributions for a selected analog.

Figure S2a shows the 4 proton configurations that were investigated. The four configurations explore three atoms that can be protonated, the carbonyl oxygen (C=O)^+^, the amide nitrogen (Amide N)^+^, and the piperidine ring. Protonation at the piperidine nitrogen results in distinct stereochemical protonation isomers (stereo-protomers). In one stereo-protomer, the proton is oriented in the same plane as the carbonyl oxygen to produce an (N-O)^+^ bridge interaction. In the other stereo-protomer, the proton is pointed in the opposite plane as the carbonyl oxygen (Piperidine-N^+^). Figures S2c-d show the relative energetics of a representative number of conformers (max 100) of these 4 protonation configurations in the gas phase as well as in water and methanol. In all media, protonation at the piperidine nitrogen results in the most energetical favored configurations. In methanol (Fig. S4d) and water (Fig. S4c), the two stereo-protomers are nearly degenerate energetically. In the gas phase (Fig S4b), the (N-O)^+^ is the most stable configuration followed by the Piperidine-N^+^ configuration. While literature suggests that the C=O^+^ protonation is present we assume that this configuration would be energetically pushed to adopt the (N-O)^+^ bridge configuration rather than maintaining an unsolvated protonated carbonyl. In fact, in all media all but one geometry of the protonated carbonyl geometries (Fig S2 b-d, pink) has relative energies of ~80 kJ/mol. Conformer 81, however, has a relative energy near zero and corresponds to the extended protonated carbonyl conformer forming a (N-O)^+^ bonding interaction. In other words, these two starting configurations, C=O^+^ and (N-O)^+^ are on the same potential energy surface, likely without a large barrier to interconversion. From this energy analysis we see that the most logical protonation configurations are the stereo-protomers, (N-O)^+^ and piperidine-N^+^, as they are the most stable in all media. Conversely protonation at the amide nitrogen is energetically unfavorable in all media.

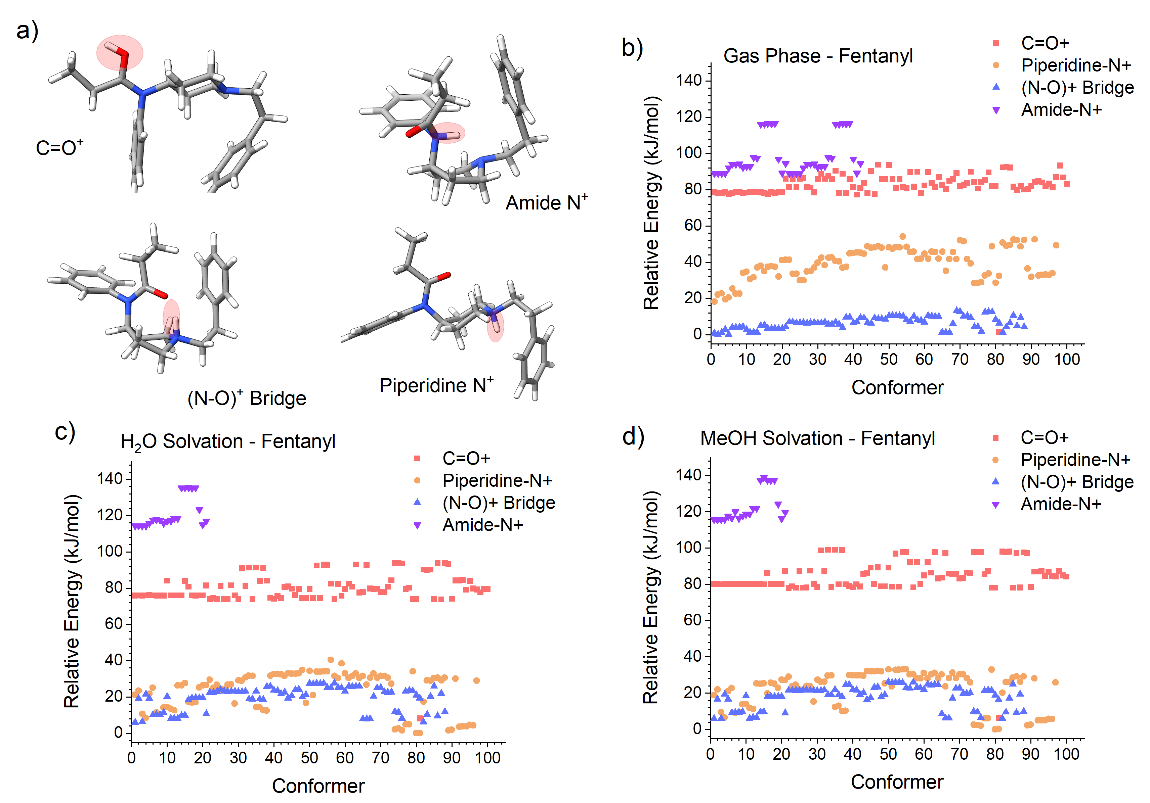

**Supplementary** **Figure S2.** Relative energies of 4 possible protonation configurations of fentanyl (m/z 337) in dielectric equivalent to methanol (a), the gas phase (b), and water (c). (d) 3-D structures of the various protonation site configurations considered.

Figure S3 plots the distribution of CCS values obtained for fentanyl (m/z 337) when modeled in the (N-O)+ and piperidine-N^+^ configurations. The distributions of CCS values were converted to Gaussian distributions by finding the average and standard deviation of the CCS values across the 100 representative gas phase conformers of each configuration. Experimental CCS values for the two mobility distributions of fentanyl were obtained from the supplementary materials of reference 27 in the main text. In this analysis, we see that the (N-O)^+^ configuration on average adopts smaller geometries compared to the piperidine-N^+^ configuration. This trend qualitatively matches the trend that is observed for experimental distributions which are shown with dotted lines.

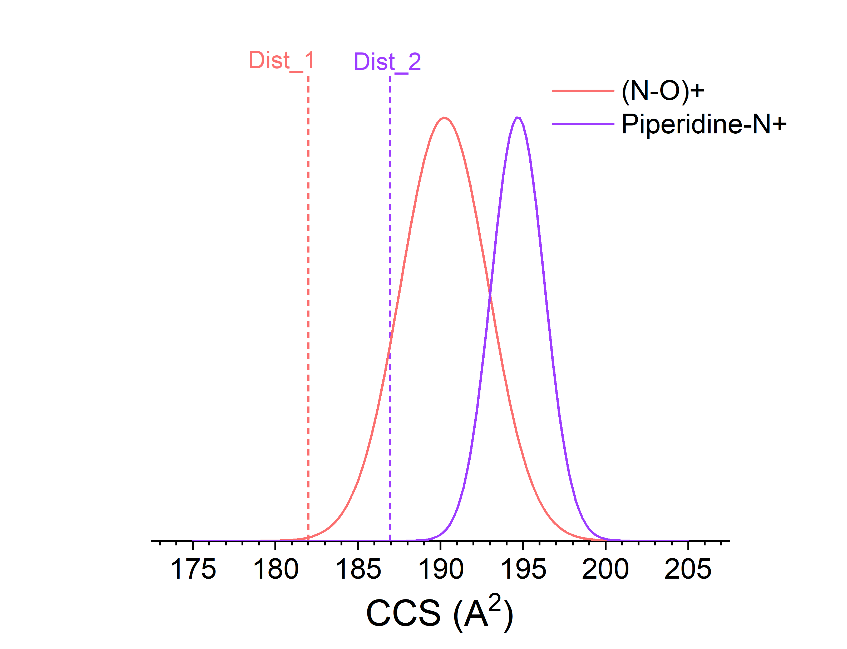

**Figure S3.** Dotted lines indicate the experimental CCS values previously measured for the two mobility distributions adopted by fentanyl (m/z 337). Solid Gaussian distributions show the distribution of CCS values adopted by 100 representative conformers of the (N-O)^+^ and piperidine-N^+^ stereoprotomers.

The top panels of Figure S4(a and b) show the predicted MS^2^ spectra of fentanyl when modeled in the (N-O)^+^ and piperidine-N^+^ configurations. MS^2^ predictions were made using QCxMS which is a molecular dynamics simulation program. While this approach is more expensive relative to ML based predictions, it allows for differences in fragmentation patterns due to protonation site to be resolved. The bottom panels in Figure S4(a and b) show the normalized fragmentation patterns obtained for the two mobility distributions of fentanyl (m/z 337). Data was obtained from the supplementary material of reference 27 in the main text. The butterfly plot in Fig S4 (a) shows that the predicted spectrum of the (N-O)^+^ configuration strongly agrees to the fragmentation pattern associated with the smaller CCS distribution of fentanyl. Specifically, m/z 188 is the most abundant fragment followed by m/z 105. The butterfly plot in Fig S4 (b) shows relatively good agreement between the predicted MS^2^ spectrum of the piperidine-N^+^ configuration and the MS^2^ spectrum of the larger CCS distribution of fentanyl. In the experimental spectrum, the most abundant fragment ion is m/z 216; this is not explicitly captured in the predicted spectrum. We do, however, see that the calculated spectrum captures the reduction in the abundance of m/z 188 as well as the presence of the other fragment ion at m/z 134.

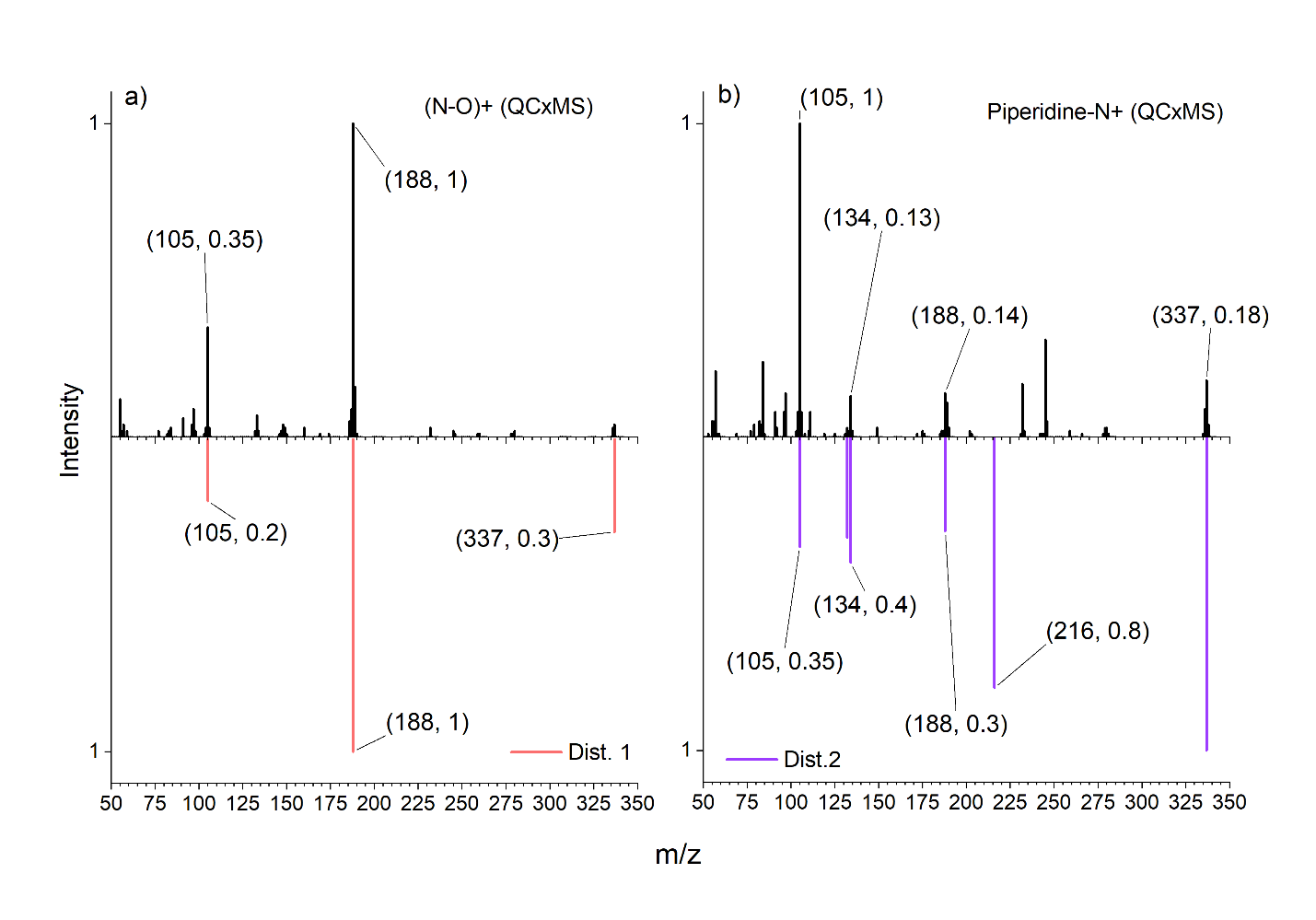

**Figure S4.** Butterfly plot comparing the calculated MS^2^ spectra of the (N-O)^+^ and the piperidine-N^+^ configurations to the experimentally observed MS^2^ spectrum of the smaller CCS distribution (a) and larger CCS distribution (b) of fentanyl (m/z 337).

Figure S5(a-g) shows the IR spectrum collected for distribution 2 of the flagged m/z values (nominal m/z) 405, 391, 385, 375, 369, 365, 351 compared to the IR spectrum of distribution 1 (Fig. S5(h-j)) for m/z values 375, 405, and 391. The highest frequency transition between 1300-1800 cm^-1^ expected for fentanyl is the carbonyl C=O stretch around ~1690 cm^-1^, corresponding to its ‘free’ position, i.e. not engaged in a hydrogen bond. This frequency is expected to shift lower as the carbonyl oxygen engages in a hydrogen bond. To further assess if the second distribution is associated with the piperidine-N^+^ configuration, we compared the highest frequency stretch recorded for distribution 2 to that of distribution 1. We only show the IR spectra collected for distribution 1 that have a SNR high enough to discern this transition. All IR spectra recorded for distribution 2 shows the highest frequency transition is around ~1660 cm^-1^. By contrast, the highest frequency transition observed for IR spectra collected on distribution 1 is between 1610-1630 cm^-1^. IR spectra collected for distribution 2 compared to 1 for the same m/z values show that the carbonyl transition is consistently shifted by 30-40 cm^-1^ (Fig. S5 (d vs. h), (a vs. i), and (b vs. j)). The shift to lower frequency is strong indication that the first distribution is associated with the (N-O)^+^ bridging interaction where the carbonyl engages in a strong hydrogen bond with the amide nitrogen and further confirms the larger distribution is associated with the piperidine-N^+^ configuration.

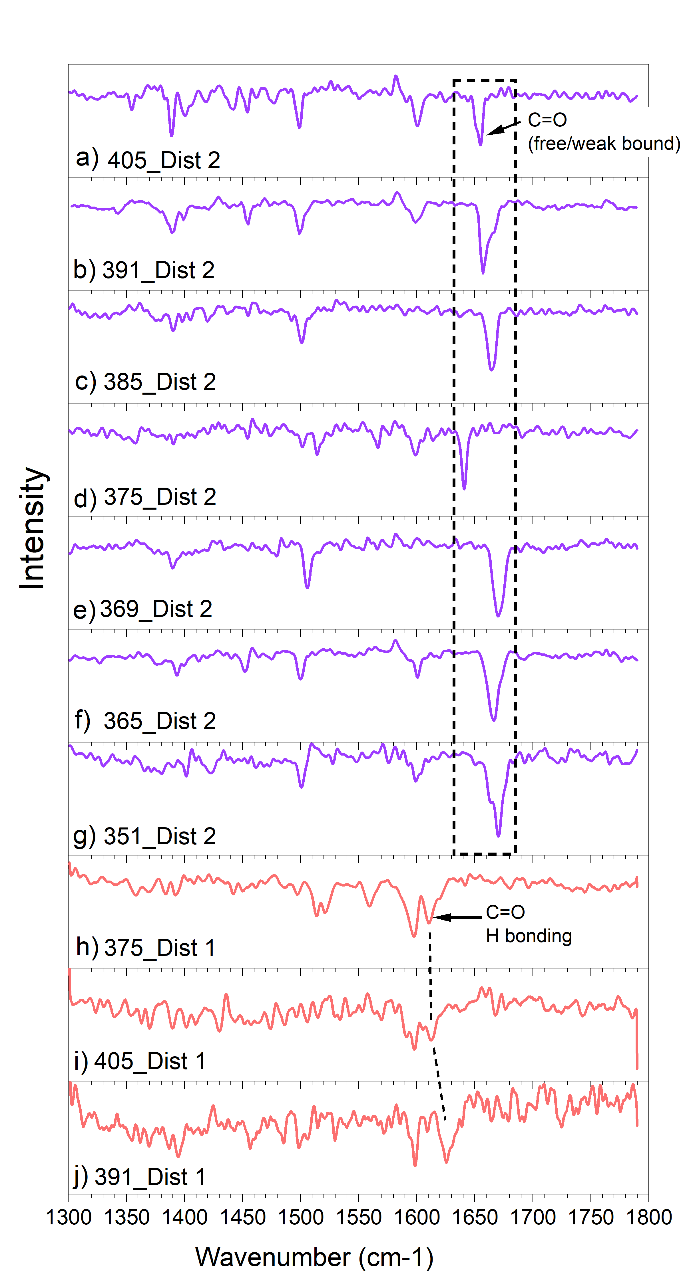

**Figure S5.** IR spectra collected for the larger CCS distribution of nominal m/z values 405, 391, 385, 369, 365, 351 (a-g), and for the smaller CCS distribution of nominal m/z values 375, 405, 391 (h-j).

**Conformation of protonation site analysis after unblinding**

Figure S6 shows the IR spectrum recorded for CCS distribution 1 of m/z 375 (top) compared to the best matched IR spectrum of the TP for m/z 375 when modeled in the (N-O)^+^ bridged configuration. Computed IR spectra show the shift in the IR frequency of the C=O band to ~1600, confirming that this geometry involves strong hydrogen bonding of the C=O.

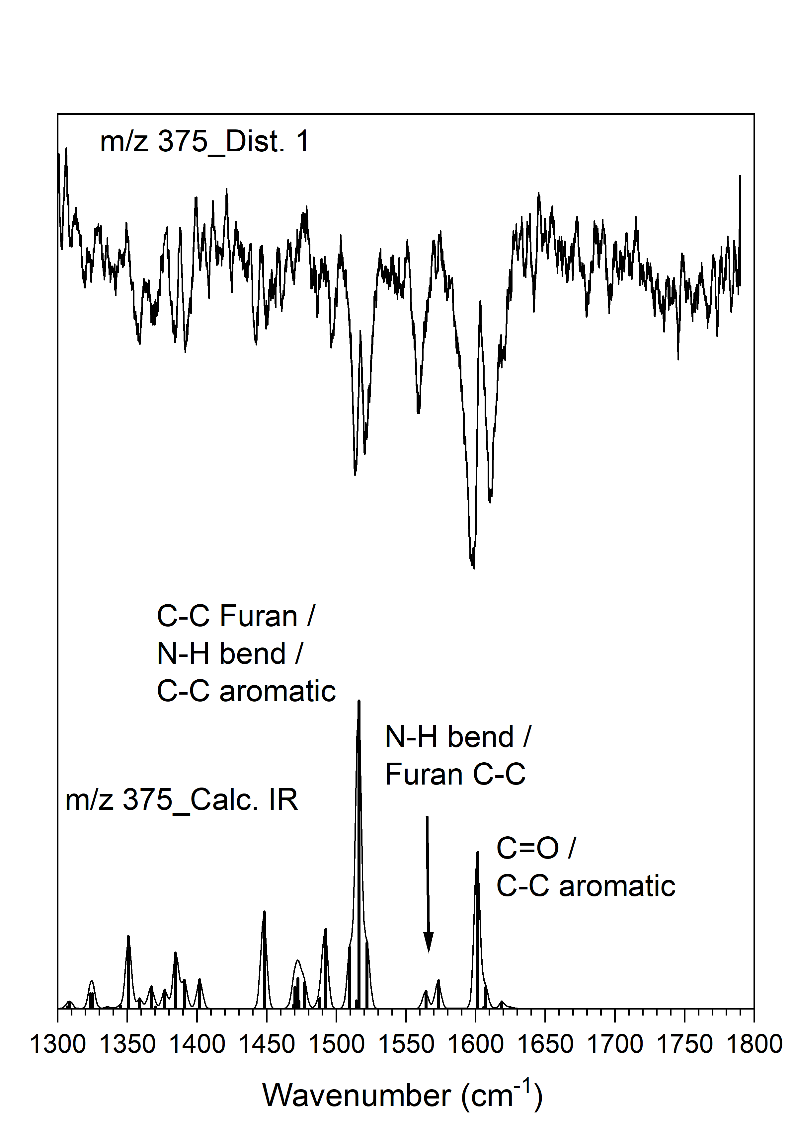

**Figure S6.** IR spectrum of m/z 375 distribution 1 (top) compared to the best matched calculated IR spectrum of the TP (bottom). In the calculated spectrum the diagnostic C=O stretch appears at ~1600 cm^-1^ rather than ~1640 cm^-1^ as observed for the calculated and experimental IR spectrum associated with distribution 2. Note that the aromatic C-C and C=O stretches in the calculated IR spectrum shown here are not resolved after applying a Gaussian broadening, leading to the appearance of a single band centered at ~1600 cm^-1^. Experimentally, these bands do however, appear well resolved yielding transitions at ~1600 cm^-1^ and ~1620 cm^-1^.

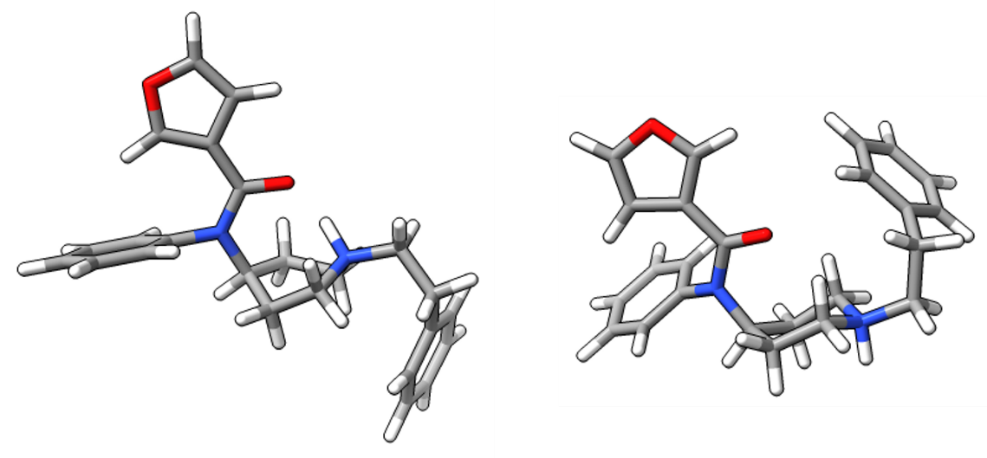

**Figure S7.** Conformers of the TP analog for m/z 375 that showed the best match in the IR domain of when compared to the IR of distribution 1 (left) and distribution 2 (right). The two conformers show the distinct protonation site stereoisomerism that gets trapped in the gas phase leading to two distinct CCS distributions and IR spectra.

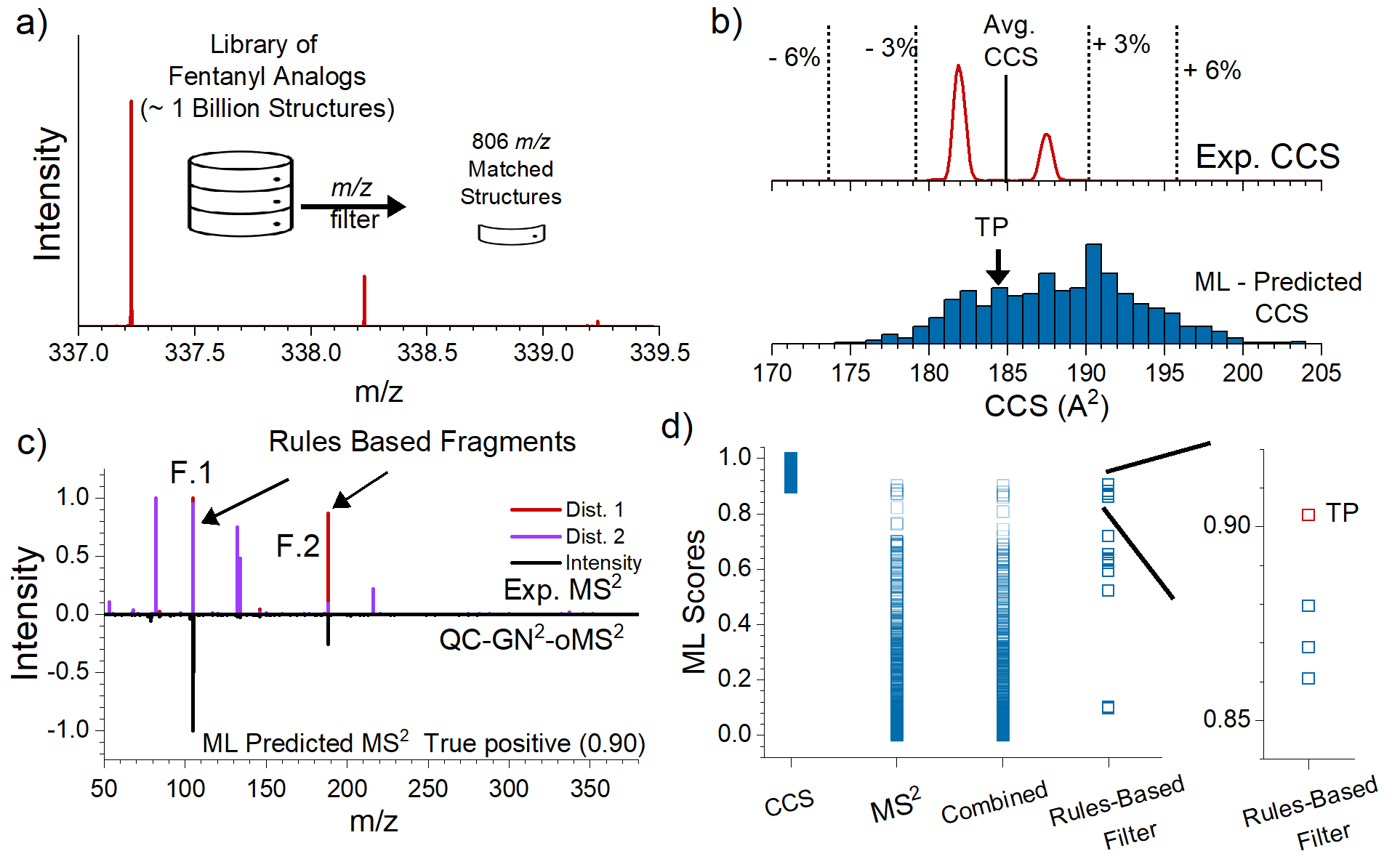

**Supplementary** **Figure S8. Down selection and identification process for flagged m/z 337.** a) HRMS measurement and mass matching against target library to reduce potential candidates from > 1 bn to 806. (b) Experimentally measured CCS distributions (top) and histogram of ML calculated CCS values for all mass matched analogs (bottom). CCS bin containing the TP is identified with a black arrow. (c) Butterfly plot comparing the experimentally measured MS^2^ spectra recorded for both CCS distributions (top) to the ML predicted MS^2^ spectrum of the TP analog (bottom). (d) Summary of 806 ML scores in the CCS and MS^2^ domains, the combined scores, as well as the scores for the remaining candidates after the rules-based analysis. The right panel shows a zoomed region containing the scores of the candidates that fall within the top 10 %; the TP is plotted in red. An IR spectrum was not collected for this m/z, and the identification was made after the ML and rules-based analysis. The TP was ranked #1.

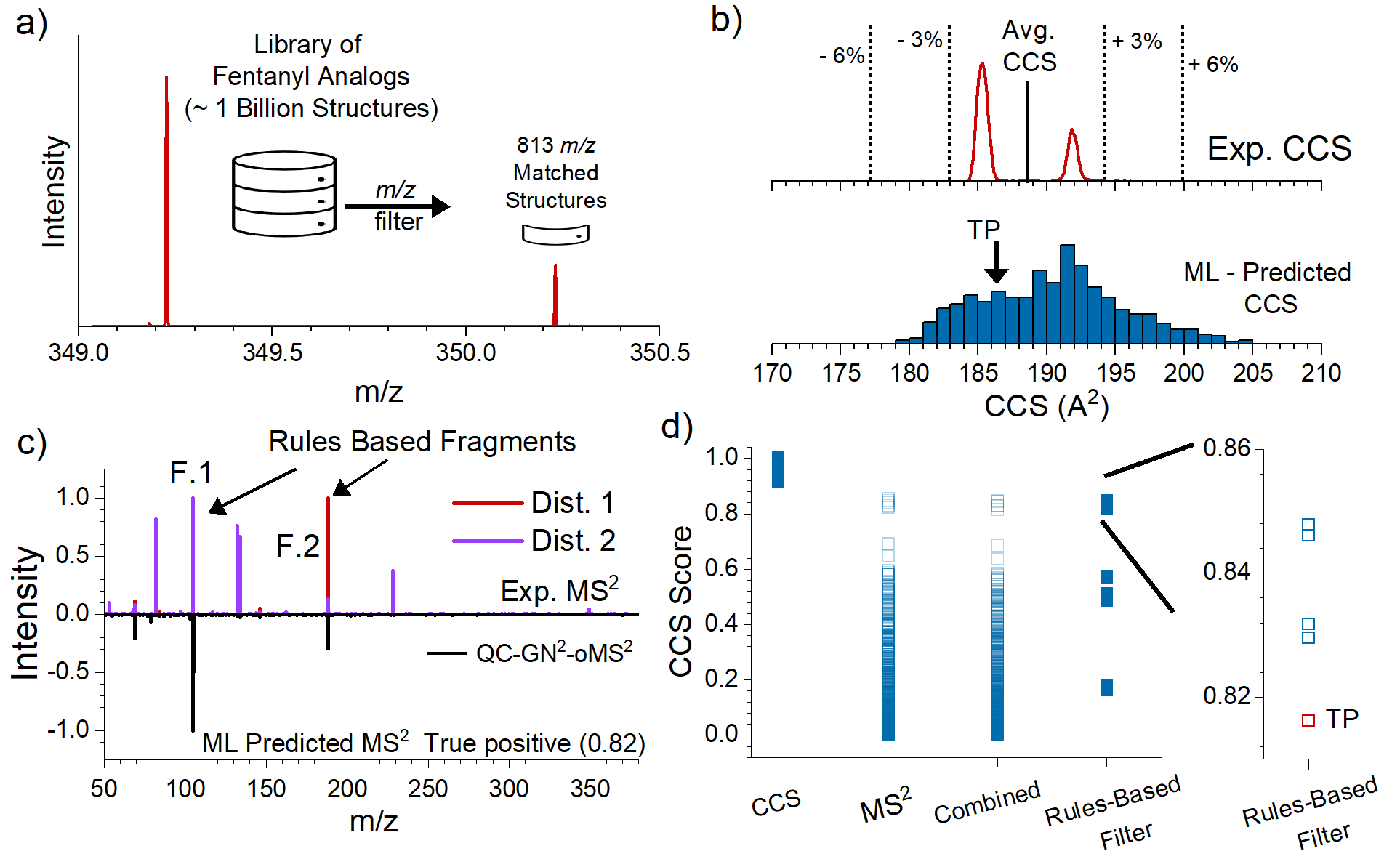

**Supplementary** **Figure S9. Down selection and identification process for flagged m/z 349.** a) HRMS measurement and mass matching against target library to reduce potential candidates from > 1 bn to 813. (b) Experimentally measured CCS distributions (top) and histogram of ML calculated CCS values for all mass matched analogs (bottom). CCS bin containing the TP is identified with a black arrow. (c) Butterfly plot comparing the experimentally measured MS^2^ spectra recorded for both CCS distributions (top) to the ML predicted MS^2^ spectrum of the TP analog (bottom). (d) Summary of 806 ML scores in the CCS and MS^2^ domains, the combined scores, as well as the scores for the remaining candidates after the rules-based analysis. The right panel shows a zoomed region containing the scores of the candidates that fall within the top 10 %, the TP is plotted in red. An IR spectrum was not collected for this m/z, and the identification was made after the ML and rules-based analysis. The TP was ranked #4.

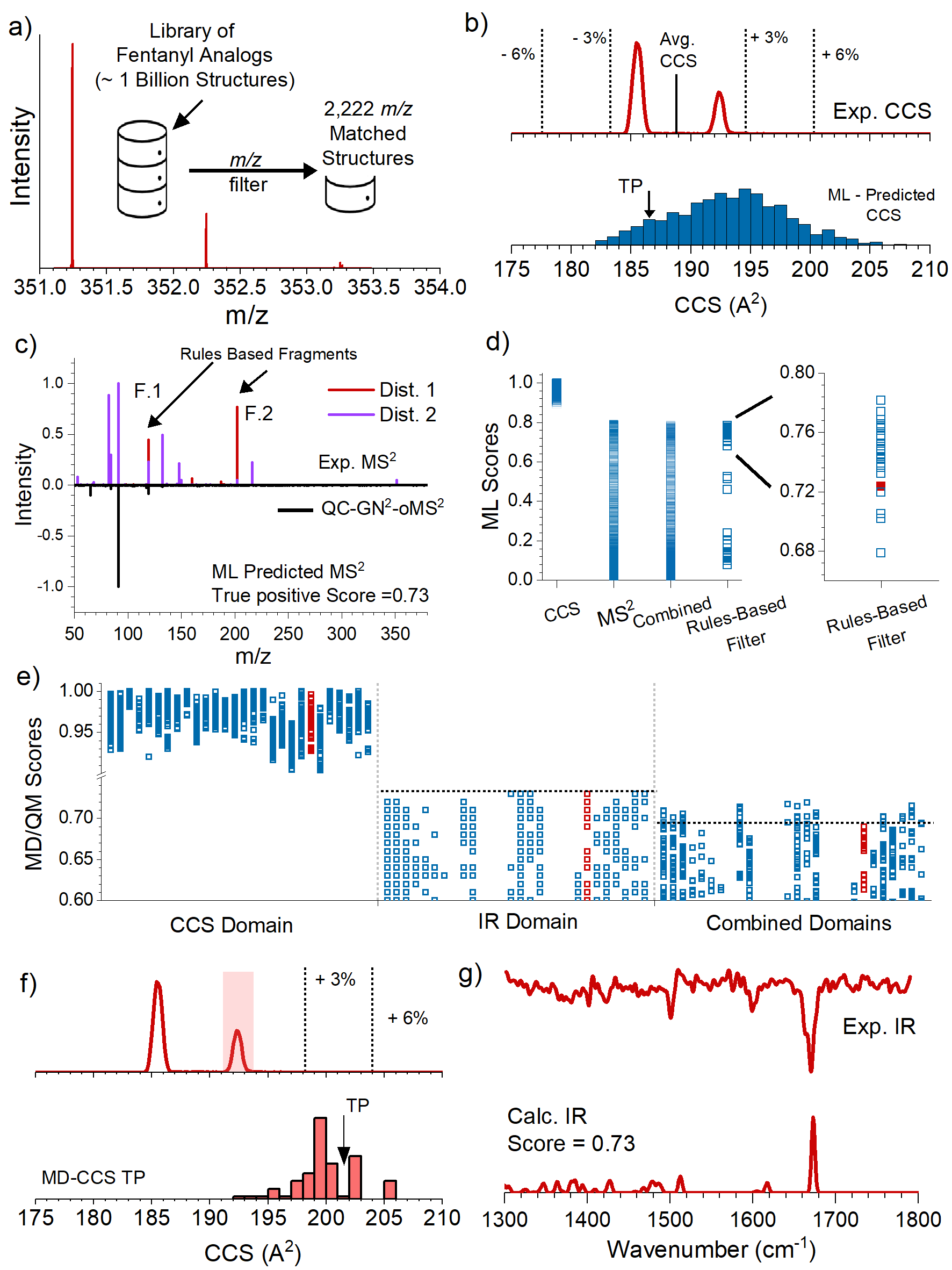

**Supplementary** **Figure S10. Down selection and identification process for flagged m/z 351.** a) HRMS measurement and mass matching against target library to reduce potential candidates from > 1 bn to 2,222. (b) Experimentally measured CCS distributions (top) and histogram of ML calculated CCS values for all mass matched analogs (bottom). CCS bin containing the TP is identified with a black arrow. (c) Butterfly plot comparing the experimentally measured MS^2^ spectra recorded for both CCS distributions (top) to the ML predicted MS^2^ spectrum of the TP analog (bottom). (d) Summary of 2,222 ML scores in the CCS and MS^2^ domains, the combined scores, as well as the scores for the remaining candidates after the rules-based analysis. The right panel shows a zoomed region containing the scores of the candidates that fall within the top 10% (27 total), the TP is plotted in red. (e) MD/QM scores of ~100 representative conformers for each top scoring analog from stage 2. Scores are split into 3 sections, CCS, IR, and the combined score. Scores associated with the TP analog are plotted in red. Dashed lines are provided as a guide for the reader to compare relative scores of true negative isomers to the true positive. (f) Comparison of the experimental CCS distribution (top) to the MD calculated CCS values for the TP analog. The shaded distribution in the experimental CCS plot is used for comparison at the MD and QM level (see main text for details). The black arrow shows the bin that contains the conformer with the highest combined CCS and IR score. (g) Comparison of the experimental (top) and best matching calculated IR spectrum (bottom) of the TP analog. The TP was ranked #12.

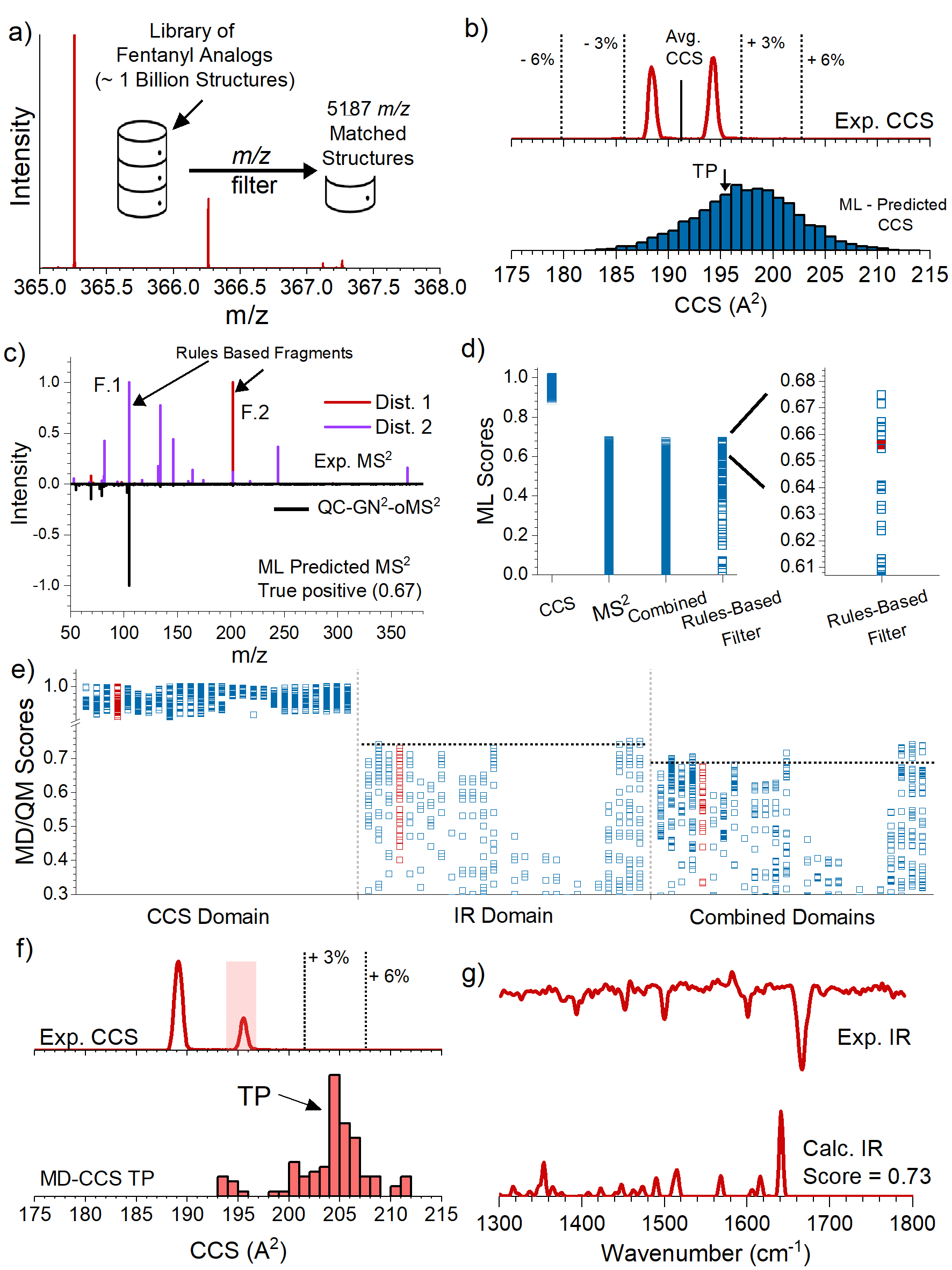

**Supplementary** **Figure S11. Down selection and identification process for flagged m/z 365.** a) HRMS measurement and mass matching against target library to reduce potential candidates from > 1 bn to 5,187. (b) Experimentally measured CCS distributions (top) and histogram of ML calculated CCS values for all mass matched analogs (bottom). CCS bin containing the TP is identified with a black arrow. (c) Butterfly plot comparing the experimentally measured MS^2^ spectra recorded for both CCS distributions (top) to the ML predicted MS^2^ spectrum of the TP analog (bottom). (d) Summary of 5,187 ML scores in the CCS and MS^2^ domains, the combined scores, as well as the scores for the remaining candidates after the rules-based analysis. The right panel shows a zoomed region containing the scores of the candidates that fall within the top 10% (26 total), the TP is plotted in red. (e) MD/QM scores of ~100 representative conformers for each top scoring analog from stage 2. Scores are split into 3 sections, CCS, IR, and the combined score. Scores associated with the TP analog are plotted in red. Dashed lines are provided as a guide for the reader to compare relative scores of true negative isomers to the true positive. (f) Comparison of the experimental CCS distribution (top) to the MD calculated CCS values for the TP analog. The shaded distribution in the experimental CCS plot is used for comparison at the MD and QM level (see main text for details). The black arrow shows the bin that contains the conformer with the highest combined CCS and IR score. (g) Comparison of the experimental (top) and best matching calculated IR spectrum (bottom) of the TP analog. The TP was ranked #7.

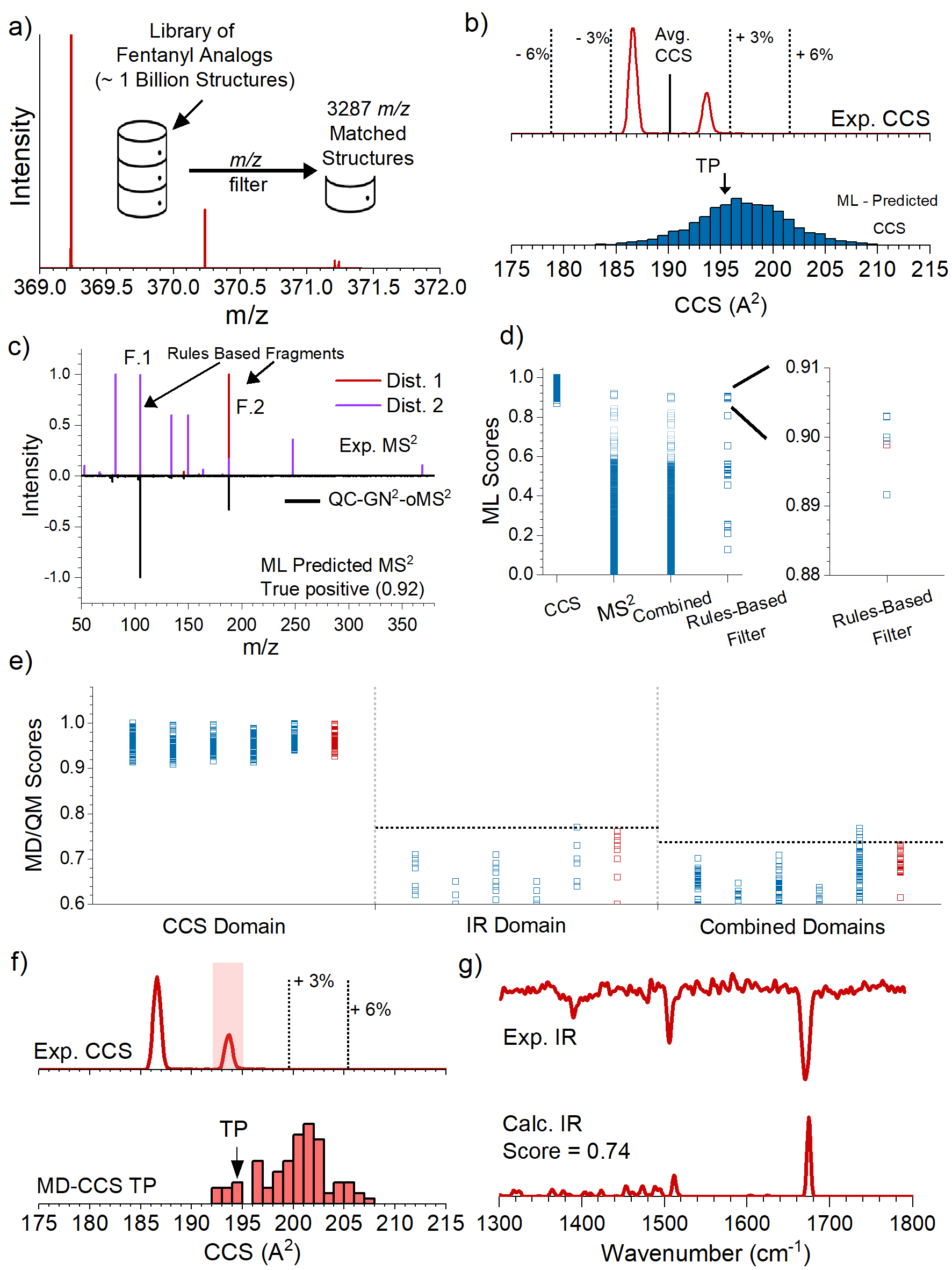

**Supplementary** **Figure S12. Down selection and identification process for flagged m/z 369.** a) HRMS measurement and mass matching against target library to reduce potential candidates from > 1 bn to 3,287. (b) Experimentally measured CCS distributions (top) and histogram of ML calculated CCS values for all mass matched analogs (bottom). CCS bin containing the TP is identified with a black arrow. (c) Butterfly plot comparing the experimentally measured MS^2^ spectra recorded for both CCS distributions (top) to the ML predicted MS^2^ spectrum of the TP analog (bottom). (d) Summary of 3,287 ML scores in the CCS and MS^2^ domains, the combined scores, as well as the scores for the remaining candidates after the rules-based analysis. The right panel shows a zoomed region containing the scores of the candidates that fall within the top 10% (6 total), the TP is plotted in red. (e) MD/QM scores of ~100 representative conformers for each top scoring analog from stage 2. Scores are split into 3 sections, CCS, IR, and the combined score. Scores associated with the TP analog are plotted in red. Dashed lines are provided as a guide for the reader to compare relative scores of true negative isomers to the true positive. (f) Comparison of the experimental CCS distribution (top) to the MD calculated CCS values for the TP analog. The shaded distribution in the experimental CCS plot is used for comparison at the MD and QM level (see main text for details). The black arrow shows the bin that contains the conformer with the highest combined CCS and IR score. (g) Comparison of the experimental (top) and best matching calculated IR spectrum (bottom) of the TP analog. The TP was ranked #2.

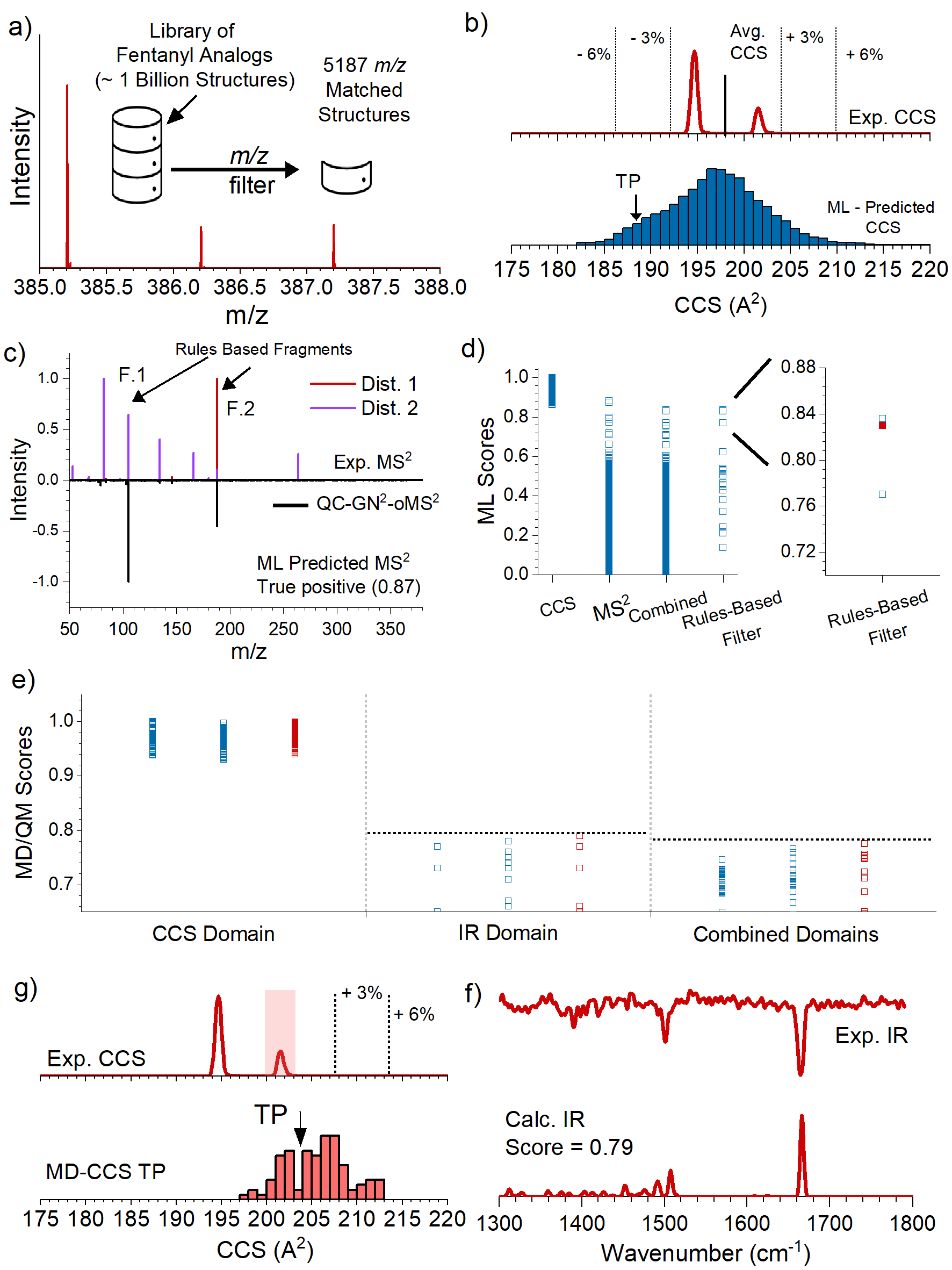

**Supplementary** **Figure S13. Down selection and identification process for flagged m/z 385.** a) HRMS measurement and mass matching against target library to reduce potential candidates from > 1 bn to 5,187. (b) Experimentally measured CCS distributions (top) and histogram of ML calculated CCS values for all mass matched analogs (bottom). CCS bin containing the TP is identified with a black arrow. (c) Butterfly plot comparing the experimentally measured MS^2^ spectra recorded for both CCS distributions (top) to the ML predicted MS^2^ spectrum of the TP analog (bottom). (d) Summary of 5,187 ML scores in the CCS and MS^2^ domains, the combined scores, as well as the scores for the remaining candidates after the rules-based analysis. The right panel shows a zoomed region containing the scores of the candidates that fall within the top 10% (3 total), the TP is plotted in red. (e) MD/QM scores of ~100 representative conformers for each top scoring analog from stage 2. Scores are split into 3 sections, CCS, IR, and the combined score. Scores associated with the TP analog are plotted in red. Dashed lines are provided as a guide for the reader to compare relative scores of true negative isomers to the true positive. (f) Comparison of the experimental CCS distribution (top) to the MD calculated CCS values for the TP analog. The shaded distribution in the experimental CCS plot is used for comparison at the MD and QM level (see main text for details). The black arrow shows the bin that contains the conformer with the highest combined CCS and IR score. (g) Comparison of the experimental (top) and best matching calculated IR spectrum (bottom) of the TP analog. The TP was ranked #1.

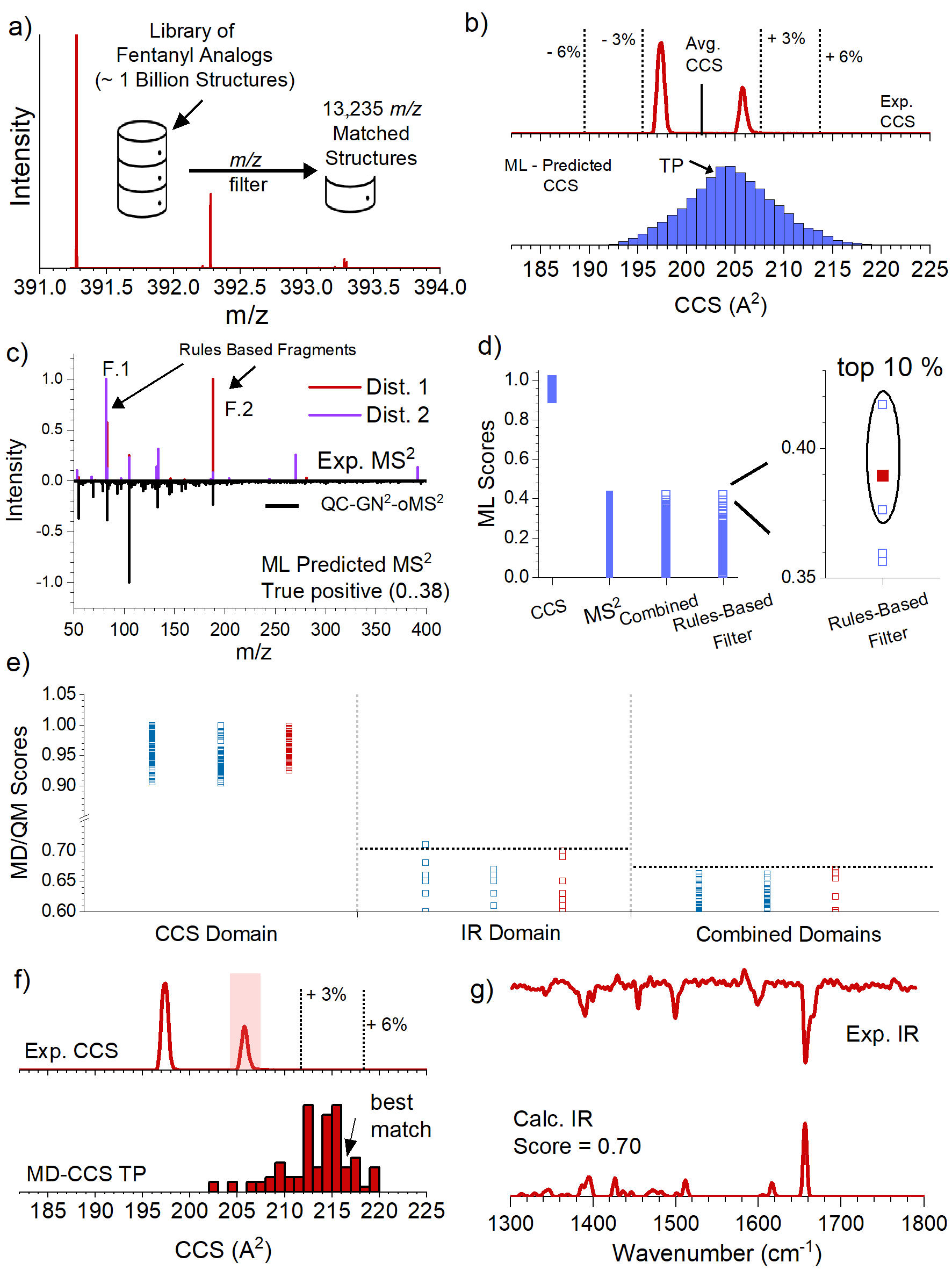

**Supplementary** **Figure S14. Down selection and identification process for flagged m/z 391.** a) HRMS measurement and mass matching against target library to reduce potential candidates from > 1 bn to 13,235. (b) Experimentally measured CCS distributions (top) and histogram of ML calculated CCS values for all mass matched analogs (bottom). CCS bin containing the TP is identified with a black arrow. (c) Butterfly plot comparing the experimentally measured MS^2^ spectra recorded for both CCS distributions (top) to the ML predicted MS^2^ spectrum of the TP analog (bottom). (d) Summary of 13,235 ML scores in the CCS and MS^2^ domains, the combined scores, as well as the scores for the remaining candidates after the rules-based analysis. The right panel shows a zoomed region containing the scores of the candidates that fall within the top 10% (3 total), the TP is plotted in red. (e) MD/QM scores of ~100 representative conformers for each top scoring analog from stage 2. Scores are split into 3 sections, CCS, IR, and the combined score. Scores associated with the TP analog are plotted in red. Dashed lines are provided as a guide for the reader to compare relative scores of true negative isomers to the true positive. (f) Comparison of the experimental CCS distribution (top) to the MD calculated CCS values for the TP analog. The shaded distribution in the experimental CCS plot is used for comparison at the MD and QM level (see main text for details). The black arrow shows the bin that contains the conformer with the highest combined CCS and IR score. (g) Comparison of the experimental (top) and best matching calculated IR spectrum (bottom) of the TP analog. The TP was ranked #1.

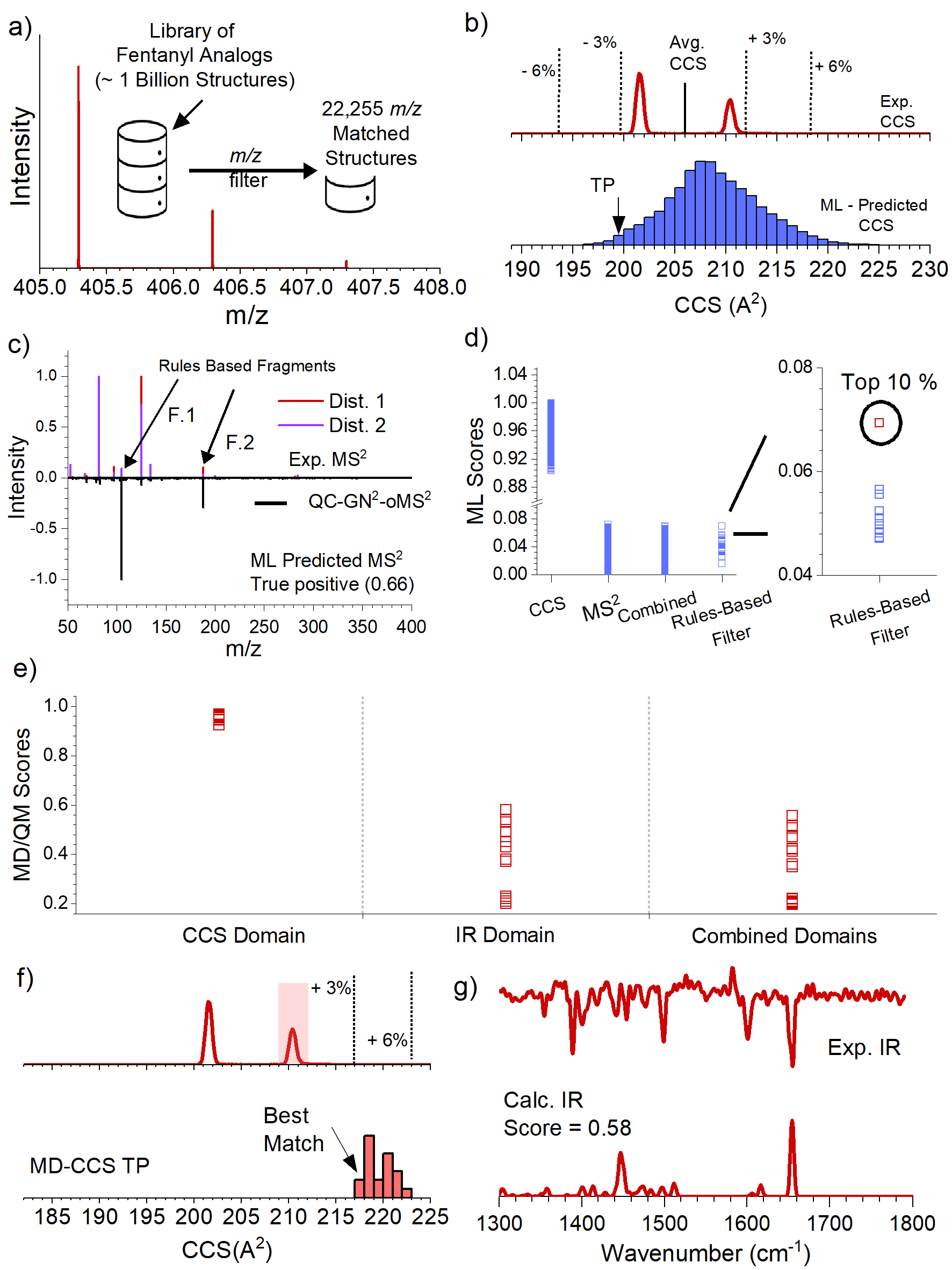

**Supplementary** **Figure S15. Down selection and identification process for flagged m/z 405.** a) HRMS measurement and mass matching against target library to reduce potential candidates from > 1 bn to 22,255. (b) Experimentally measured CCS distributions (top) and histogram of ML calculated CCS values for all mass matched analogs (bottom). CCS bin containing the TP is identified with a black arrow. (c) Butterfly plot comparing the experimentally measured MS^2^ spectra recorded for both CCS distributions (top) to the ML predicted MS^2^ spectrum of the TP analog (bottom). (d) Summary of 22,255 ML scores in the CCS and MS^2^ domains, the combined scores, as well as the scores for the remaining candidates after the rules-based analysis. The right panel shows a zoomed region containing the scores of the candidates that fall within the top 10% (1 total), the TP is plotted in red. (e) MD/QM scores of ~100 representative conformers for each top scoring analog from stage 2. Scores are split into 3 sections, CCS, IR, and the combined score. Scores associated with the TP analog are plotted in red. Dashed lines are provided as a guide for the reader to compare relative scores of true negative isomers to the true positive. (f) Comparison of the experimental CCS distribution (top) to the MD calculated CCS values for the TP analog. The shaded distribution in the experimental CCS plot is used for comparison at the MD and QM level (see main text for details). The black arrow shows the bin that contains the conformer with the highest combined CCS and IR score. (g) Comparison of the experimental (top) and best matching calculated IR spectrum (bottom) of the TP analog. The TP was ranked #1.

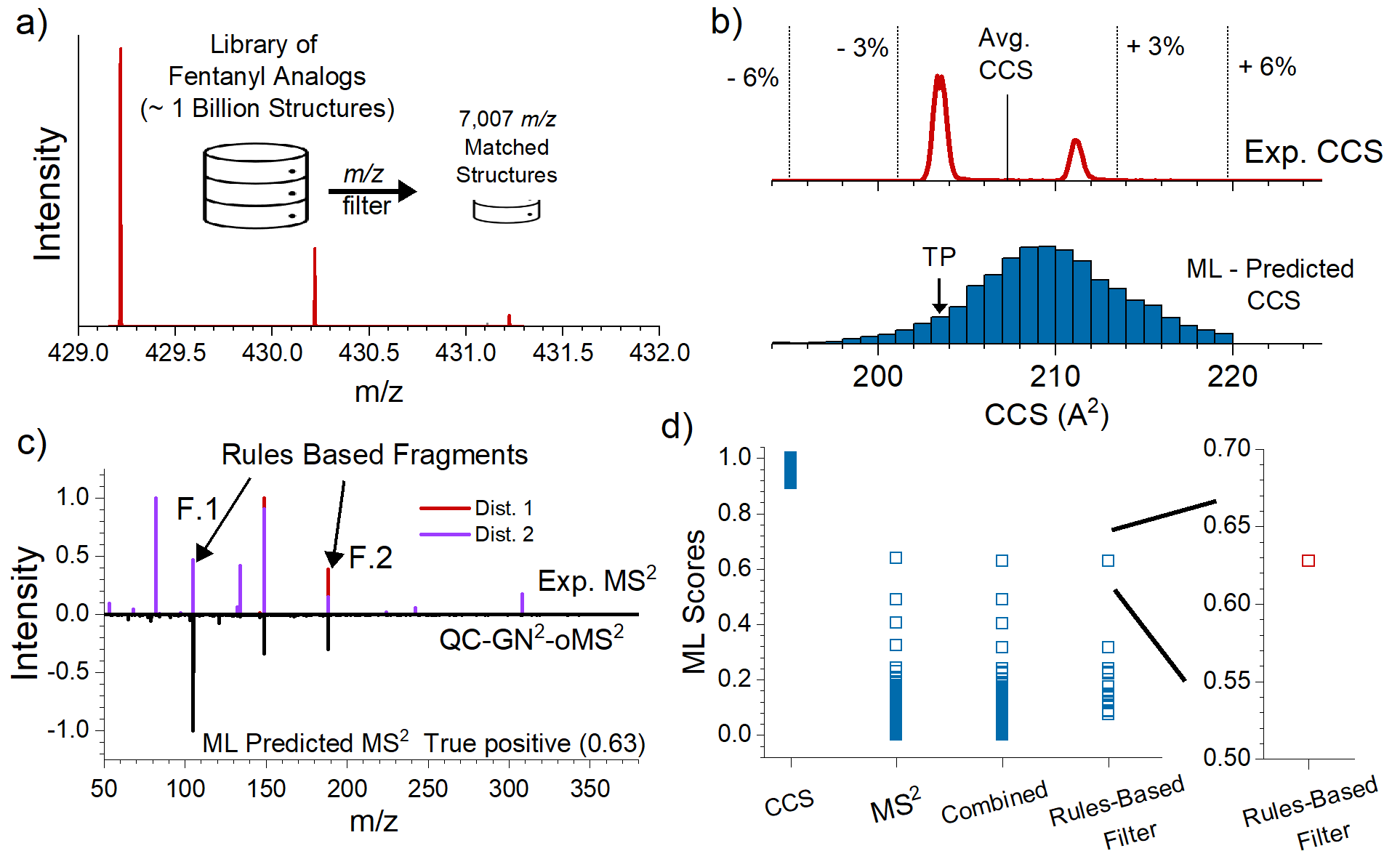

**Supplementary** **Figure S16. Down selection and identification process for flagged m/z 429.** a) HRMS measurement and mass matching against target library to reduce potential candidates from > 1 bn to 7,007. (b) Experimentally measured CCS distributions (top) and histogram of ML calculated CCS values for all mass matched analogs (bottom). CCS bin containing the TP is identified with a black arrow. (c) Butterfly plot comparing the experimentally measured MS^2^ spectra recorded for both CCS distributions (top) to the ML predicted MS^2^ spectrum of the TP analog (bottom). (d) Summary of 7,007 ML scores in the CCS and MS^2^ domains, the combined scores, as well as the scores for the remaining candidates after the rules-based analysis. The right panel shows a zoomed region containing the scores of the candidates that fall within the top 10%, the TP is plotted in red. An IR spectrum was not collected for this m/z, and the identification was made after the ML and rules-based analysis. The TP was ranked #1.

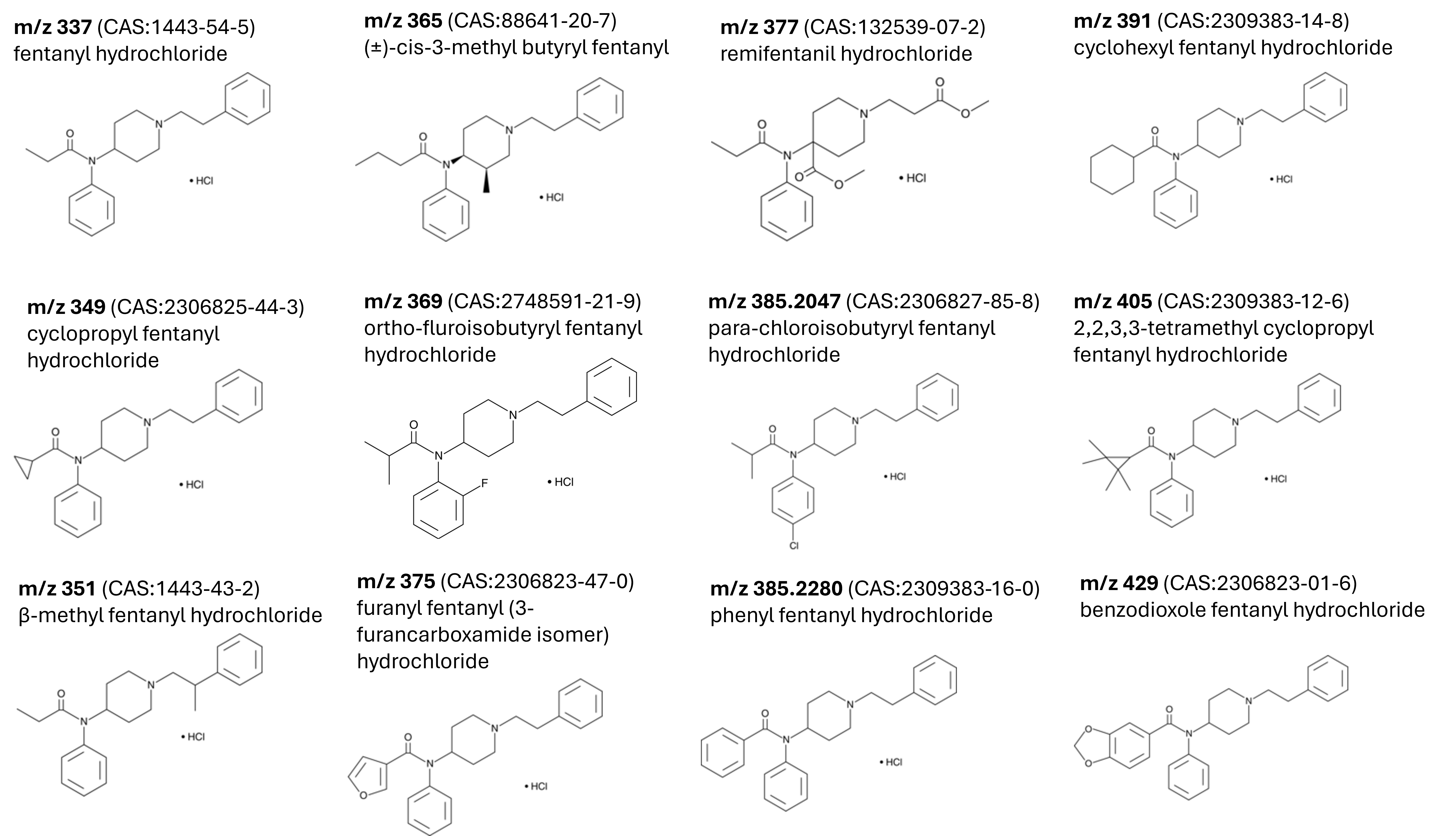

**Supplementary Figure S17.** 2-D structures of 12 fentanyl analogs used in the blinded study. Analogs were probed as protonated ions but are received as hydrochloride salts from the vendor.

**Unflagged Fentanyl Analogs**

Remifentanil did not display two distributions in the mobility domain, which was a requirement for a *m/z* to be flagged, and was not investigated as a result. The mass spectrum and CCS distribution for remifentanil is shown in Supplementary Figure S18. Phenyl fentanyl was not flagged as it was not fully resolved from para-chloroisobutyryl fentanyl (m/z 385.2047) in the SLIM-cryoIR-ToF. These analogs have *m/z* differences of ~60 ppm. Supplementary Figure S17 shows the ability to resolve between these two analogs in *m/z* and CCS domain on the SLIM-Orbitrap. The lower resolution in the *m/z* domain of the SLIM-cryoIR-ToF however, prevented these two analogs from being mass resolved rather a single peak is observed at *m/z* 385.21, Supplementary Figure S17a. Further, the slightly lower resolution in the IMS domain also prevented the two analogs from being fully resolved by their mobilities under the experimental conditions used, Supplementary Figure S17b. As a result, the measured values of the SLIM-cryoIR-ToF were associated with the measured values of para-chloroisobutyryl fentanyl from the SLIM-Orbitrap, as the *m/z* and CCS values were closer in agreement. In the post analysis, we confirmed that the IR spectrum of 385.21 was largely representative of para-chloroisobutyryl fentanyl. This was done by comparing the IR spectrum associated with *m/z* 387.21 to that of *m/z* 385.21. *m/z* 387.21 is the (M+2) isotope associated with the chlorine present in para-chloroisobutyryl fentanyl. Given the larger natural abundance of ^37^Cl compared ^13^C, the IR spectrum of 387.21 is nearly exclusively representative of the para-chloroisobutyryl fentanyl. This comparison is made in Supplementary Figure S17 and shows that the IR spectrum collected for 385.21 is indeed associated with para-chloroisobutyryl fentanyl. Based on the width of the CCS distributions observed for *m/z* 385.21, we can ascertain the presence of phenyl fentanyl in the mixture; however, we did not attempt to reconstruct its IR spectrum.

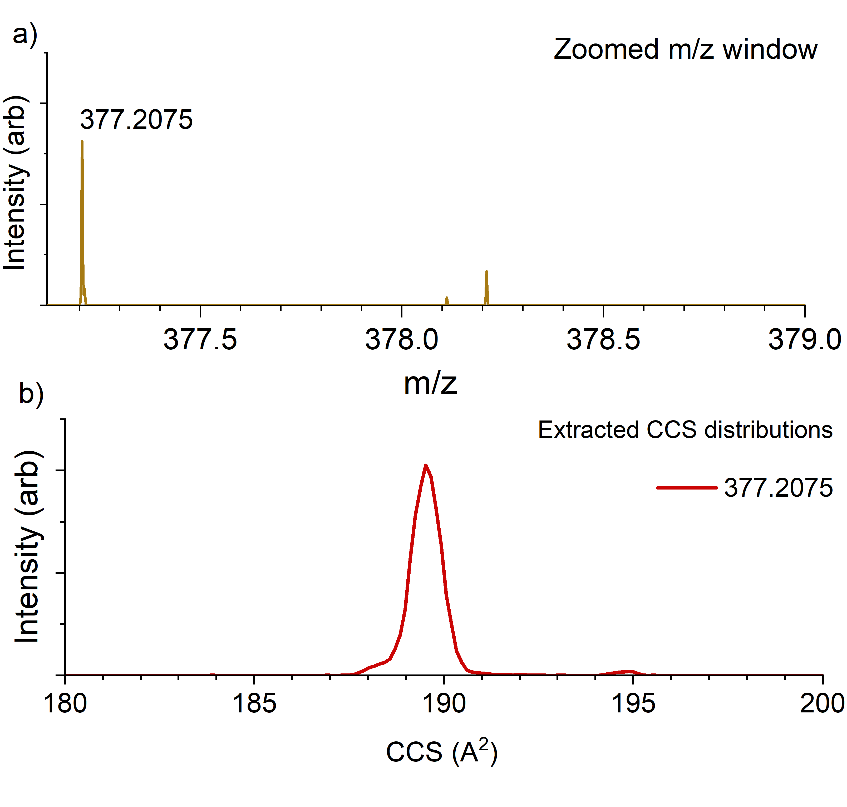

**Supplementary** **Figure S18.** (a) Zoomed m/z window around remifentanil. (b) Extracted CCS distribution of remifentanil showing 1 distribution. Data shown was collected on the SLIM-Orbitrap. The m/z corresponding to remifentanil was not flagged as a potential fentanyl analog as it did not show 2 distinct distributions in the CCS domain.

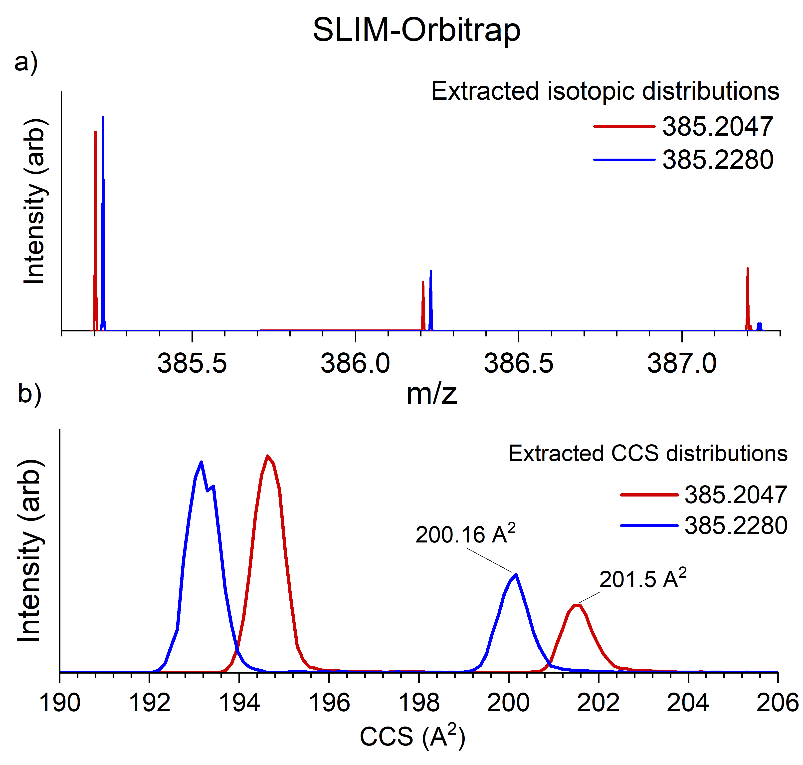

**Supplementary Figure S19.** (a) Zoomed m/z window (m/z 385.1-387.3) showing measured m/z values of para-chloroisobutyryl fentanyl (m/z 385.2047) and phenyl fentanyl (m/z 385.2280). (b) Extracted CCS values for m/z 385.2047 and 385.2280. Data was recorded on SLIM-Orbitrap. Analogs differ by 0.0233 amu (~60 ppm) and are readily resolved by m/z and CCS on the SLIM-Orbi.

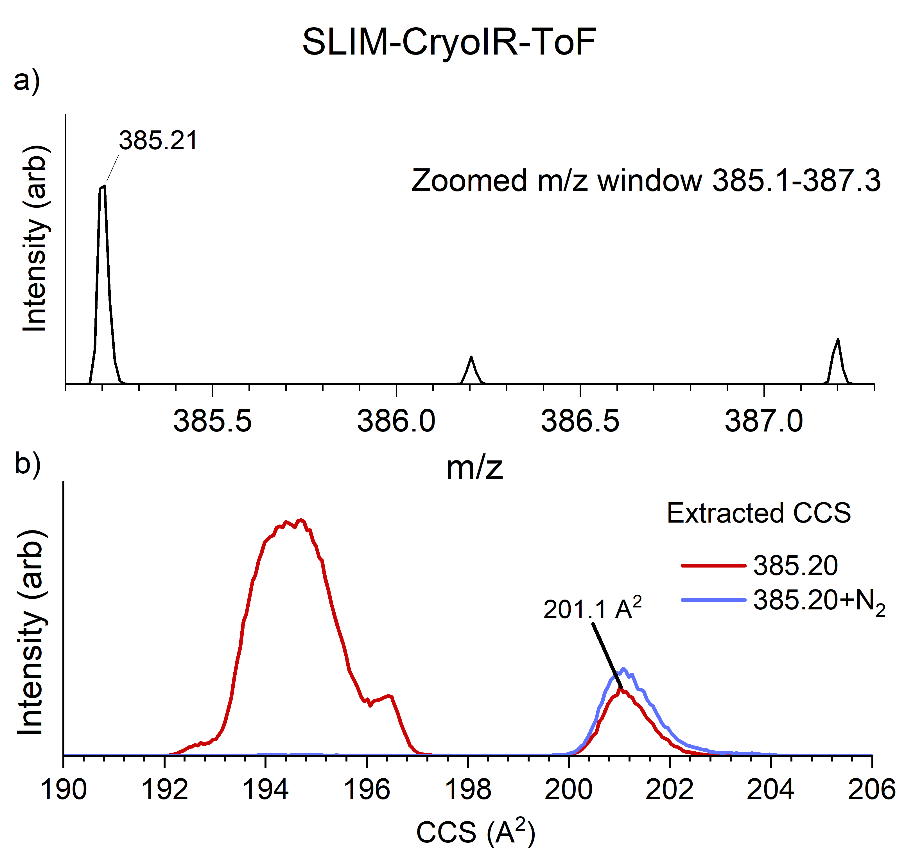

**Supplementary Figure S20.** (a) Zoomed window between m/z 385.1-387.3, showing unresolved para-chloroisobutyryl fentanyl (m/z 385.2047) and Phenyl fentanyl (m/z 385.2280). (b) Extracted CCS distribution for m/z 385.21 showing limited separation of para-chloroisobutyryl fentanyl and phenyl fentanyl due to lower IMS resolutions on the SLIM-cryoIR-ToF. Measured values for this species were linked to measured properties of m/z 385.2047 based on closer agreement in the m/z and IMS domains.

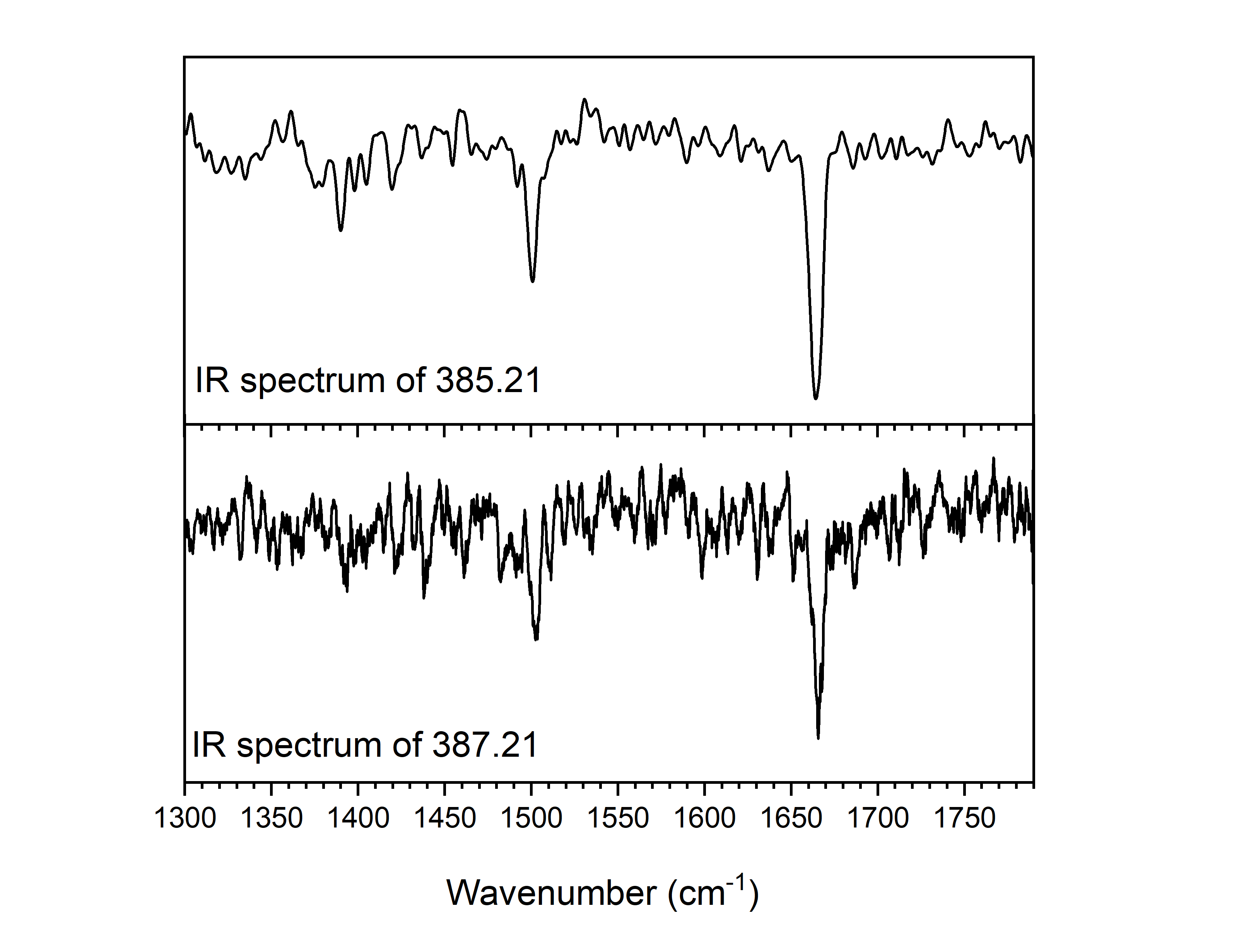

**Supplementary Figure S20.** IR spectrum recorded for m/z 385.21 and 387.21 (the ^37^Cl isotope of chloroisobutyryl fentanyl). The spectra are identical, indicating that the spectrum recorded for 385.21 is largely attributed to chloroisobutyryl fentanyl.

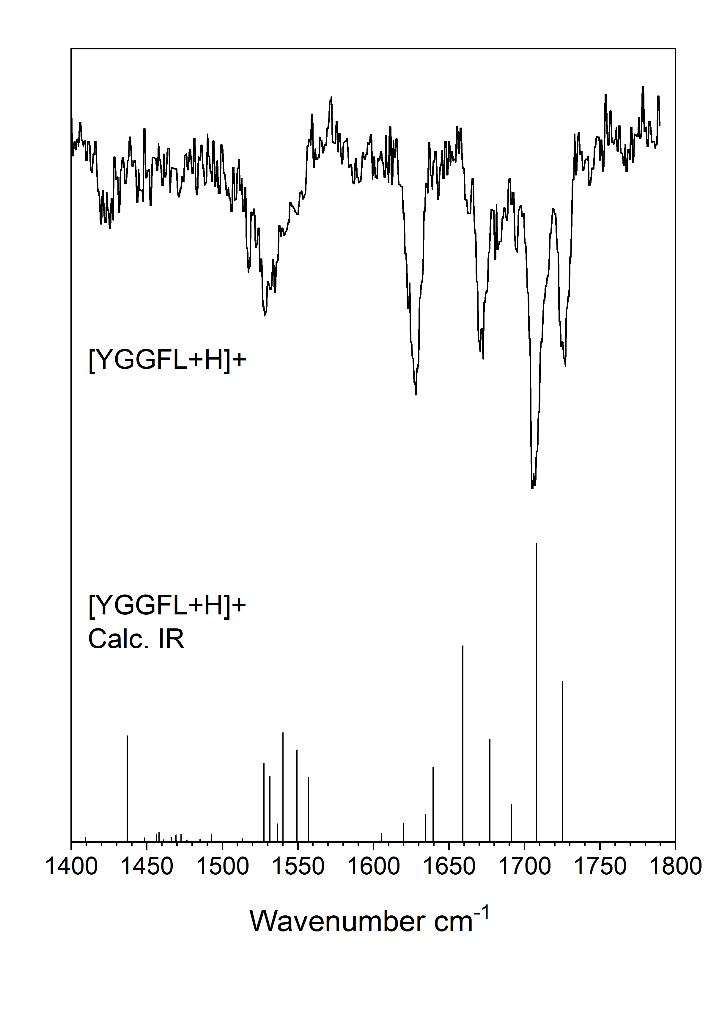

**Figure S21.** Comparison of experimental spectrum of [YGGFL+H]^+^ (top) to IR spectrum calculated using “B3LYP d3 def2-TZVP KDIIS RIJCOSX def2/J TIGHTOPT VerySlowConv TIGHTSCF FREQ defgrid2” level of theory and basis set with Orca 5.0 after scaling by 0.986.

**Crest Input:**

srun -N 1 --ntasks-per-node=1 --mpi=none crest $JOBDIR/$MYXYZ --noreftopo --chrg +1 -T 64

**Orca input for geometry and IR predictions:**

! B3LYP d3 def2-TZVP KDIIS RIJCOSX def2/J TIGHTOPT VerySlowConv TIGHTSCF FREQ defgrid2

%maxcore 4000

%base "CREST"

%pal nprocs 32

end

%scf

MaxIter 1500

convergence tight

end

%method

Z_solver Pople # Pople usually recommended. DIIS sometimes.

Z_MaxIter 300

Z_Tol 1e-5

end

* xyzfile +1 1

**XYZ coordinates of [YGGFL+H]^+^**

N	-2.58069	0.46323	-0.60593

C	-3.33352	1.60998	0.01200

C	-2.32497	2.62176	0.55766

O	-2.27598	3.74754	0.07614

C	-4.32665	1.05570	1.04051

C	-5.22502	0.01912	0.40715

C	-4.96759	-1.34689	0.57454

C	-5.75050	-2.30735	-0.05272

C	-6.81587	-1.90846	-0.86121

C	-7.08914	-0.55050	-1.03835

C	-6.29392	0.40037	-0.40727

O	-7.54770	-2.88909	-1.44448

N	-1.51122	2.20056	1.54846

C	-0.36965	3.01995	1.94184

C	0.71319	2.79388	0.89487

O	1.57466	1.92292	1.02915

N	0.59611	3.51305	-0.23767

C	1.37229	3.10079	-1.38325

C	1.09801	1.63244	-1.70249

O	-0.05090	1.17581	-1.70835

N	2.17176	0.87891	-1.98436

C	2.05401	-0.52515	-2.32618

C	1.56471	-1.38418	-1.16534

O	1.13855	-2.52296	-1.41277

C	3.38337	-1.06262	-2.89239

C	4.54899	-0.81048	-1.96732

C	4.79283	-1.65944	-0.88346

C	5.83840	-1.39877	-0.00260

C	6.65368	-0.28366	-0.19184

C	6.42753	0.56203	-1.27428

C	5.38231	0.29620	-2.15853

N	1.58038	-0.88013	0.07519

C	0.99325	-1.62025	1.19378

C	-0.51778	-1.49747	1.00274

O	-1.23625	-0.74085	1.64114

C	1.45913	-1.06310	2.53108

C	2.97620	-1.14942	2.74209

C	3.47696	-2.59467	2.70254

C	3.32634	-0.49838	4.08120

O	-1.04444	-2.18980	0.00403

H	-3.78224	0.61707	1.88075

H	-4.90249	1.89885	1.42781

H	-4.15275	-1.66683	1.21710

H	-5.56297	-3.36448	0.08109

H	-7.92345	-0.23761	-1.65623

H	-6.52507	1.45244	-0.53974

H	-8.28324	-2.52039	-1.95026

H	-3.85493	2.11831	-0.79644

H	-2.23487	-0.17091	0.13269

H	-3.21178	-0.09073	-1.19068

H	-1.76089	0.77377	-1.15748

H	-0.68204	4.06114	1.99916

H	-0.00544	2.68325	2.90830

H	-1.50061	1.22732	1.83254

H	1.07353	3.70101	-2.24213

H	2.43474	3.26086	-1.19541

H	-0.24006	4.06968	-0.36684

H	3.09833	1.24757	-1.80852

H	1.27876	-0.63156	-3.08621

H	3.55418	-0.58123	-3.85727

H	3.24733	-2.13025	-3.07126

H	4.17396	-2.53999	-0.74438

H	6.02550	-2.07276	0.82410

H	7.47098	-0.08628	0.49000

H	7.07197	1.41588	-1.44174

H	5.23485	0.93490	-3.02482

H	1.26904	-2.67069	1.08346

H	1.83498	0.09005	0.24351

H	1.13744	-0.02285	2.60916

H	0.94953	-1.62256	3.32206

H	3.47484	-0.58397	1.94716

H	4.53542	-2.63838	2.96716

H	3.36814	-3.04894	1.71436

H	2.93132	-3.21099	3.42397

H	4.40550	-0.50540	4.24540

H	2.85789	-1.04156	4.90735

H	2.98532	0.53896	4.11637

H	-0.32816	-2.61920	-0.54129
